# Uncovering structural ensembles from single particle cryo-EM data using cryoDRGN

**DOI:** 10.1101/2022.08.09.503342

**Authors:** Laurel F. Kinman, Barrett M. Powell, Ellen D. Zhong, Bonnie Berger, Joseph H. Davis

**Affiliations:** Department of Biology, Massachusetts Institute of Technology, Cambridge, MA; Computational and Systems Biology Graduate Program, Massachusetts Institute of Technology, Cambridge, MA; Computer Science and Artificial Intelligence Laboratory, Massachusetts Institute of Technology, Cambridge, MA; Department of Mathematics, Massachusetts Institute of Technology, Cambridge, MA

**Author notes:** These authors contributed equally (Laurel F. Kinman, Barrett M. Powell, Ellen D. Zhong).

## Abstract

CryoDRGN is a machine learning system for heterogeneous cryo-EM reconstruction of proteins and protein complexes from single particle cryo-EM data. Central to this approach is a deep generative model for heterogeneous cryo-EM density maps, which we empirically find effectively models both discrete and continuous forms of structural variability. Once trained, cryoDRGN is capable of generating an arbitrary number of 3D density maps, and thus interpreting the resulting ensemble is a challenge. Here, we showcase interactive and automated processing approaches for analyzing cryoDRGN results. Specifically, we detail a step-by-step protocol for analysis of the assembling 50S ribosome dataset (Davis et al., EMPIAR-10076), including preparation of inputs, network training, and visualization of the resulting ensemble of density maps. Additionally, we describe and implement methods to comprehensively analyze and interpret the distribution of volumes with the assistance of an associated atomic model. This protocol is appropriate for structural biologists familiar with processing single particle cryo-EM datasets and with moderate experience programming in Python and navigating Jupyter notebooks. It requires 3-4 days to complete.

## INTRODUCTION

Proteins and their complexes exist in dynamic equilibria: assembling, disassembling, and undergoing conformational changes. Many of these dynamics are intrinsically linked to function, yet, often, they are poorly understood on a structural level. In recent years, cryo-EM has emerged as a powerful tool for studying protein structure (Lyumkis, 2019; Serna, 2019; Wu and Lander, 2020) and the single-molecule nature of cryo-EM makes it an appealing choice for studying protein motions, as millions of individual particles sampled from an underlying energy landscape can be visualized on a single grid (Dashti et al., 2014; Haselbach et al., 2017; Dashti et al., 2020; Zhong et al., 2020; Gui et al., 2021; Punjani and Fleet, 2021b). However, studying highly heterogeneous cryo-EM datasets has proved to be a challenging computational problem, as most traditional approaches rely on extensive classification and particle averaging (Grant et al., 2018; Punjani et al., 2017; Zivanov et al., 2018) to produce approximately static structures, thereby gaining resolution while obscuring or blurring underlying structural variation (Nakane et al., 2018; Punjani and Fleet, 2021b; Zhong et al., 2020).

We have developed an approach that leverages machine learning models capable of embedding heterogeneous single particle cryo-EM images within a low-dimensional latent space and generating 3D volumes as a function of that latent embedding (Zhong et al., 2020). Our approach, named cryoDRGN, takes as inputs a particle stack and poses from a consensus 3D refinement, and uses these data to train a neural network architecture based on the variational autoencoder (VAE) (Kingma and Welling, 2013; Zhong et al., 2019). The overall architecture consists of two neural networks: an image encoder network, which assigns a latent embedding ***Z**_i_* to each particle *i*, and a volume decoder network, which reconstructs a 3D density map *_V__i_* given ***z**_i_*. The development, theoretical foundations, and limitations of this work have been described previously (Zhong et al., 2019; Zhong et al., 2020). We have applied this approach to a number of publicly-available datasets, and found that cryoDRGN can uncover rare structural states in assembling bacterial ribosomes (Sun et al., 2022) and help visualize continuous conformational changes in spliceosome complexes (Zhong et al., 2020). CryoDRGN has also helped visualize a tilting motion of radial spike proteins important in dynein motors and ciliary motility (Gui et al., 2021).

To illustrate the process of training a cryoDRGN model on a cryo-EM dataset and interpreting the resulting outputs, we present a full protocol and pipeline to analyze an assembling large ribosomal subunit dataset — EMPIAR-10076 (Davis et al., 2016), which exhibits rich compositional and conformational heterogeneity and has been previously characterized (Zhong et al., 2020; Punjani and Fleet, 2021b; Rabuck-Gibbons et al., 2021). The presented pipeline details: 1) the preparation of inputs for cryoDRGN given a particle stack and corresponding consensus reconstruction; 2) training of an initial low-resolution cryoDRGN model; 3) filtering the input particle stack based on the results of low-resolution training; 4) high-resolution cryoDRGN training; and 5) analysis, visualization, and interpretation of the resulting structural ensembles with the assistance of an atomic model (**Figure 1**).

**Figure 1.**
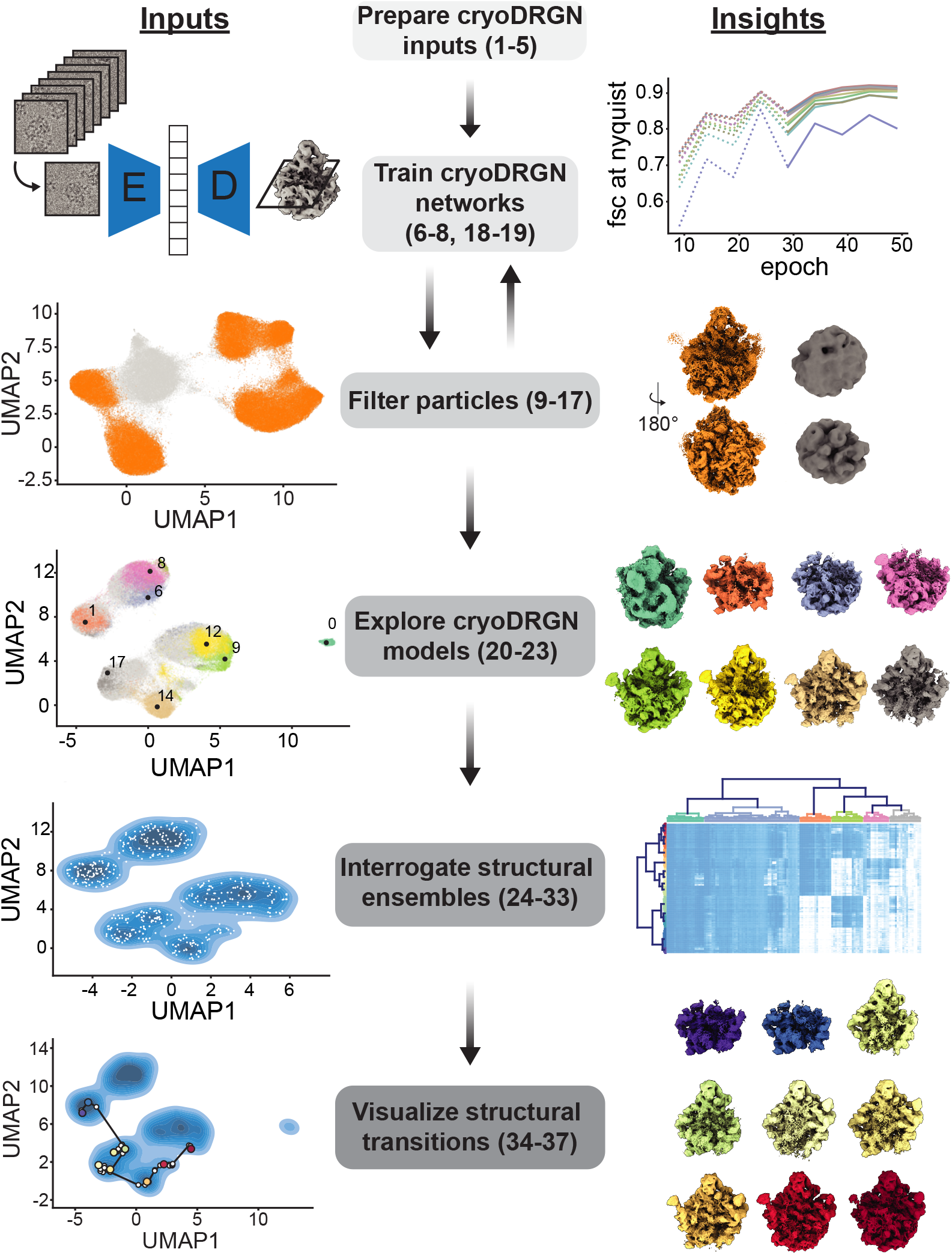
The cryoDRGN workflow. Steps (*center*) of the cryoDRGN analysis workflow are noted, with typical inputs to each step (*left*) and insights gained (*right*) illustrated. Each noted step corresponds to a subsection of the provided protocol, with numbered steps of the protocol listed.

### Comparison with other methods

Traditional approaches to handle heterogeneity rely on successive rounds of user-driven discrete 2D- and 3D-classification, which separate particles into a few independent underlying structures (Serna, 2019). Success of these approaches is strongly dependent on selection of the appropriate number of classes, which is unknown *a priori*, and initial models used for refinement, which can be a significant source of bias. Moreover, this strategy relies on a fundamental assumption that the data are well-described by a finite and identifiable number of true volumes. For datasets displaying conformational heterogeneity, in which particles exist in states sampled along one or more continuous trajectories, this assumption does not hold. Even for datasets that show large degrees of discrete compositional heterogeneity, questions still remain about how and when to stop classifying, and how robust results are to classification parameters (Rabuck-Gibbons et al., 2021).

More recently, alternatives to traditional global 3D classification, including multi-body refinement (Nakane et al., 2018) and focused classification (von Loeffelholz et al., 2017), have been developed. However, these approaches are inherently limited by the assumption that the structural heterogeneity can be decomposed into a small number of rigid bodies and that the user can identify and define these rigid bodies using masks. In contrast, a number of tools for modeling continuous heterogeneity have also emerged, including principal component analysis based approaches such as cryoSPARC’s 3D variability analysis (Punjani and Fleet, 2021b), 2 neural network based approaches like cryoDRGN (Zhong et al., 2020), 3DFlex (Punjani and Fleet, 2021a), and e2gmm (Ludtke and Chen, 2021) aimed at generating heterogeneous ensembles of 3D-density maps, and analogous methods for directly inferring ensembles of atomic models (Rosenbaum et al., 2021; Zhong et al., 2021). 3D variability analysis models heterogeneity as a linear combination of eigenvolumes, and is thus limited in its ability to model complex, nonlinear motions, whereas 3DFlex learns a single underlying structure and a set of continuous deformations of this structure and may therefore be challenged by discrete heterogeneity caused by either large cooperative movements, or by compositional variation within a complex.

CryoDRGN has broad applicability for modeling complex ensembles containing both continuous and discrete heterogeneity, with the ability to generate an arbitrary number of maps from the imaged ensemble. We have found that cryoDRGN is sufficiently powerful to model non-linear continuous motions and discrete changes in complex composition, yet, unlike many of the aforementioned methods, does not require strong structural priors like the number of expected classes or specification of rigid domains that are expected to undergo conformational changes. Here we provide a protocol detailing how cryoDRGN can be applied to an exemplary heterogeneous dataset and describe additional recently developed tools to aid in analyzing and interpreting the resulting structural ensembles.

### Overview of the procedure

This protocol was developed through comprehensive experimentation and analyses of a variety of cryo-EM datasets across a wide variety of systems. The protocol presents our best practices in real application settings.

#### Preparing cryoDRGN inputs

Within the cryo-EM single particle reconstruction pipeline, cryoDRGN is applied between the steps of traditional 3D reconstruction and model-building (**Figure 1**). As inputs, cryoDRGN requires a stack of extracted single particles and their corresponding poses and CTF parameters, which are derived from a traditional consensus 3D reconstruction in which the heterogeneous particles are aligned in the same reference frame to a single volume. In general, one should observe well-defined secondary structure in portions of the refined volume as indicative of accurately posed images before initiating cryoDRGN training. Notably, we have found forgoing stringent particle filtering at this stage often expands the range of heterogeneity cryoDRGN learns. To improve resolution, poses and per-particle CTF parameters should be optimized, optionally through the use of non-uniform refinement (Punjani et al., 2020) or Bayesian polishing (Zivanov et al., 2019).

CryoDRGN’s required inputs can be generated by many single particle reconstruction packages, and we provide preprocessing tools to convert from cryoSPARC (Punjani et al., 2017) and RELION (Scheres, 2012) output formats. During this preprocessing stage we recommend users downsample the particle stack to a lower resolution to facilitate rapid initial network training for dataset filtering. Additionally, we back-project the downsampled particle stack using the cryoDRGN-parsed inputs and compare with the refined volume to confirm that the inputs have been correctly prepared (**Extended Data Figure 1**).

#### Training cryoDRGN networks

A cryoDRGN model is trained by iterating through the dataset of particle images and updating neural network parameters with stochastic gradient descent on the loss function described below. One epoch of such training entails passing all particles through the encoder and decoder networks once. The mean squared error between each input image and the corresponding image reconstructed by the decoder network is used to estimate a *“reconstruction loss”* that is used in conjunction with a *“regularization loss”* on the latent embeddings to iteratively update the network parameters (**Figure 2A**). At the end of every iteration the updated parameters and latent space embedding for each particle are saved as `weights.[epoch].pkl` and `z.[epoch].pkl`, respectively. Thus, the output directory following 50 epochs of training will contain 50 network weights files, 50 per-particle latent embedding z files, a `config.pkl` file containing the input parameters and settings used, and a `run.log` file containing information about the run.

**Figure 2.**
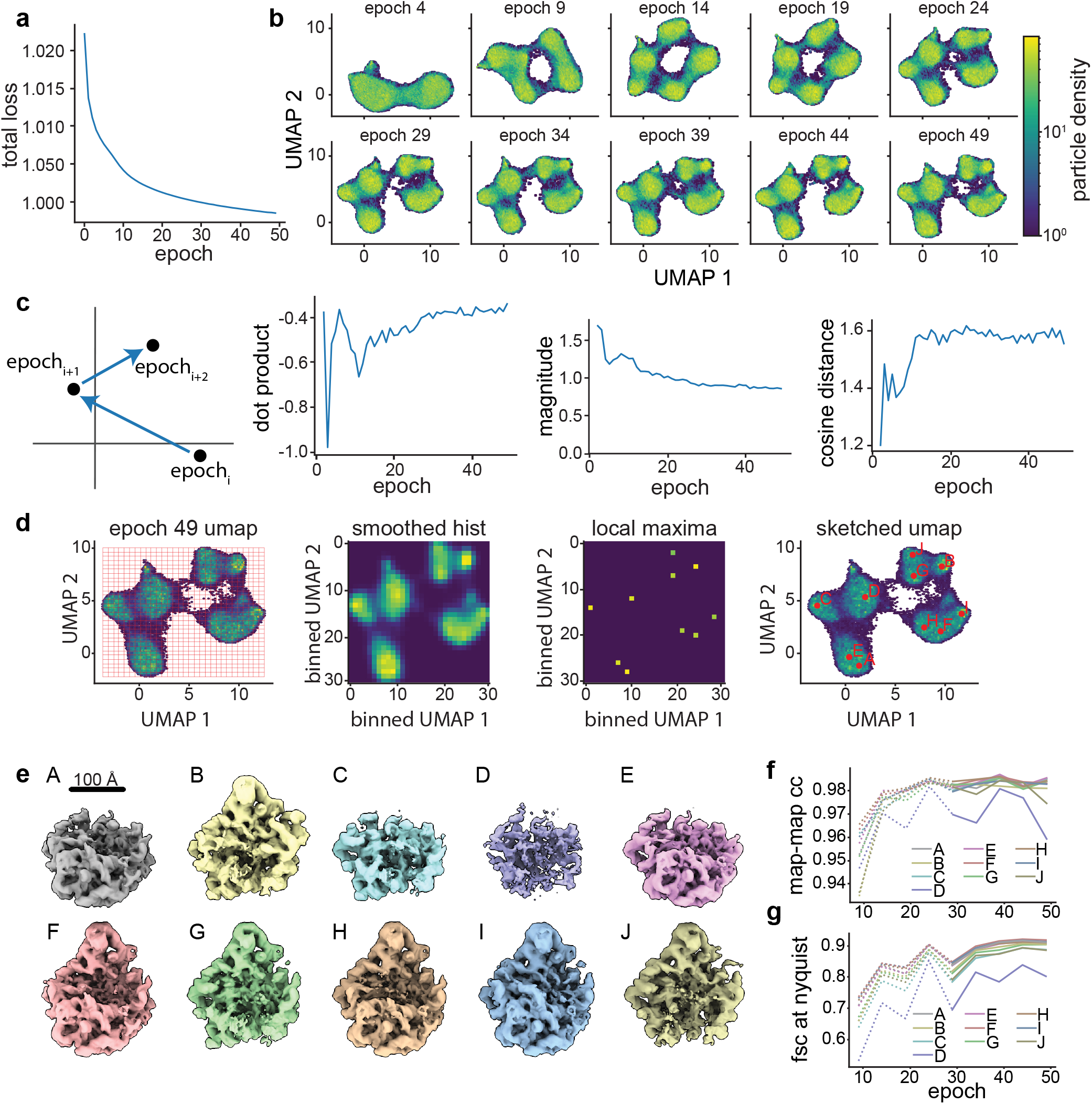
Training and assessing convergence of cryoDRGN networks. **a)** Representative plot of total loss at each epoch. Decreasing loss reflects gains in neural network performance. **b)** Representative density heatmaps of the particle embeddings at noted epochs of training. In each density heatmap, UMAP was used to embed each 8-D latent distribution in a 2-D space. Note the shape of the resulting heatmap stabilizes in later epochs, consistent with encoder network convergence. **c)** Illustration of a hypothetical particle’s embedding in successive epochs of training. Difference vectors between successive epochs are colored blue (*left).* Such vectors’ dot product, magnitude, and cosine distance are computed, and the median value at each epoch is shown (*right).* The asymptotic behavior of these curves is consistent with encoder network convergence. **d)** Identification of representative latent embeddings via the *“UMAP local maxima method”* (Glossary). **e)** Volumes generated by the decoder network at the local maxima positions (A-J) defined in d. Note the diversity of low-resolution structures. **f)** Map-to-map correlation and **g)** FSC at Nyquist frequency calculated between volumes generated from local maxima identified as defined in d at five epoch intervals. Epochs for which the encoder network was not assessed to have converged are noted with dotted lines.

Training a neural network can be computationally expensive. We have implemented GPU parallelization and accelerated mixed-precision training which can lead to considerably faster network training, particularly for large network architectures and image sizes. However, care must be taken to ensure that training has converged as training dynamics are altered when using GPU parallelization. In this protocol, we use a single GPU and mixed-precision training.

We also describe intuitive heuristics to judge when neural network training has converged and is unlikely to benefit from additional epochs of training (**Box 1, Figure 2, Extended Data Figures 2,4,5**). For simplicity, we independently assess the convergence of the encoder and decoder networks and focus our assessment on particles likely to be of interest. Here, we define particles of interest as those in well-populated neighborhoods of latent space, as these neighborhoods are, by definition, well supported by the data. First, we examine encoder convergence, and determine the training epoch after which particles of interest only minimally change their relative positions in latent space. Convergence is monitored epoch-to-epoch by visual comparison of UMAP embeddings (**Figure 2B**), and by characterizing particle movement in latent space during training (**Figure 2C**). Second, we monitor decoder convergence by assessing the training epoch after which density maps corresponding to particles of interest no longer change. Here, convergence is monitored by generating volumes from a consistent set of particles of interest and comparing map-to-map correlation coefficients and FSC curves between epochs (**Figure 2D-G, Extended Data Figures 2,4,5**). We provide a dedicated script within cryoDRGN that automatically calculates and plots these criteria. Once training of the encoder and decoder networks has converged, any subsequent epoch can be used for filtering or analysis, with the caveat that we occasionally observe overfitting to noise in epochs well beyond training convergence. Thus, we recommend examining volumes for signs of increased noise, streaking artifacts, or other pathologies in the density maps. If such pathologies are observed, an earlier epoch should be analyzed.

##### Box 1: Convergence Analysis

Here, we describe several heuristic metrics to assess convergence of cryoDRGN network training. Each metric queries convergence of different elements of the network (*i.e.* the encoder, the decoder, or the entire network). Although alternative heuristics exist, we have found that these metrics are useful in judging when cryoDRGN networks have been sufficiently trained across a variety of datasets. The motivation, implementation, and example interpretation of each convergence metric are detailed below.

###### Total network loss

This metric is the loss function guiding network learning during training. Total loss per epoch is expected to decrease as the network trains. Smooth asymptotic behavior is indicative of stable network training.

###### UMAP latent embeddings

Upon convergence of the encoder network we expect the distribution of latent embeddings to be insensitive to further training. To visualize high dimensional latent distributions, we calculate UMAP embeddings of the latent at set intervals during training. Note that UMAP is subject to artifacts like rotation, mirroring, or inconsistent mapping of particles on cluster boundaries. When applied to datasets producing a featured latent space, the important criteria are that the number, size, and relative distribution of clusters remains constant. For datasets with less featured UMAP embeddings, locally monitoring dense regions within UMAP clusters or relying on alternative metrics can be useful.

###### Latent embedding shifts

This metric examines the *“movement”* of particles through latent space during training, with the expectation that upon encoder network convergence, such movements will be small and randomly directed within local minima. Movement is monitored by the size (magnitude) and consistency of direction (dot product and cosine distance) of a given particle’s motion over epochs of training. Specifically, we consider the n-dimensional vectors connecting its latent embedding in epoch_i_, to epoch_i+1_, and in epoch_i+1_ to epoch_i+2_. The magnitude of each vector, as well as the dot product and cosine distance of this pair of vectors, are calculated and the median values for these three parameters across all particles are plotted per epoch. Similar to the total loss plot, an elbow and subsequent stabilization in each of these plots is consistent with convergence. Less featured latent spaces can result in more *“noise”* in these plots; in such cases, a rolling average of these values can be used.

###### Correlation of generated volumes

This approach assesses the convergence by examining whether volumes sequentially generated from related positions in latent space stabilize during training. These positions are calculated as the on-data median latent values of particles in well-supported clusters identified using the *“UMAP local maxima method”* (**Glossary**). The median latent encoding of each cluster’s particles is updated, and a corresponding volume generated, every five epochs. Volumes generated in this way should trend towards high correlation with the previously generated volume during convergence, as particles map to increasingly consistent regions of latent space and the decoder produces increasingly consistent corresponding volumes. Stabilization is measured by map-to-map real-space correlation and map-to-map FSC. For this dataset, which produces structures whose resolutions are Nyquist-limited, we find in addition to examining FSC at all spatial frequencies, specifically visualizing the increasing correlation at the Nyquist limit is informative.

In general, strict cutoffs for convergence are difficult to identify. These heuristics are intended to be used in a holistic fashion when assessing convergence. Typically, additional training beyond convergence provides diminishing returns while increasing the likelihood of overtraining artifacts as described above.

#### Particle filtering

Image heterogeneity within a given cryo-EM dataset can result from true structural heterogeneity of the particle of interest or from the presence of contaminants such as ice or edge artifacts. To achieve high-quality reconstructions in traditional cryo-EM processing workflows, these contaminants are often removed by iterative rounds of 2D- or 3D-classification (Cheng et al., 2015). We have observed that the latent embeddings produced by cryoDRGN can distinguish between true particles and contaminants, and thus represent a powerful alternative method to filter particle stacks (Zhong et al., 2020). Because training time scales with both the number of particles and the box size, and because the presence of contaminants consumes representation capacity in the neural networks, we recommend using an initial round of low-resolution training to eliminate contaminants before proceeding to high-resolution training. Here, we describe one round of particle filtering (**Figure 3, Extended Data Figure 3**); however, when working with other datasets, users may find it useful to iterate through multiple rounds of particle filtering.

**Figure 3.**
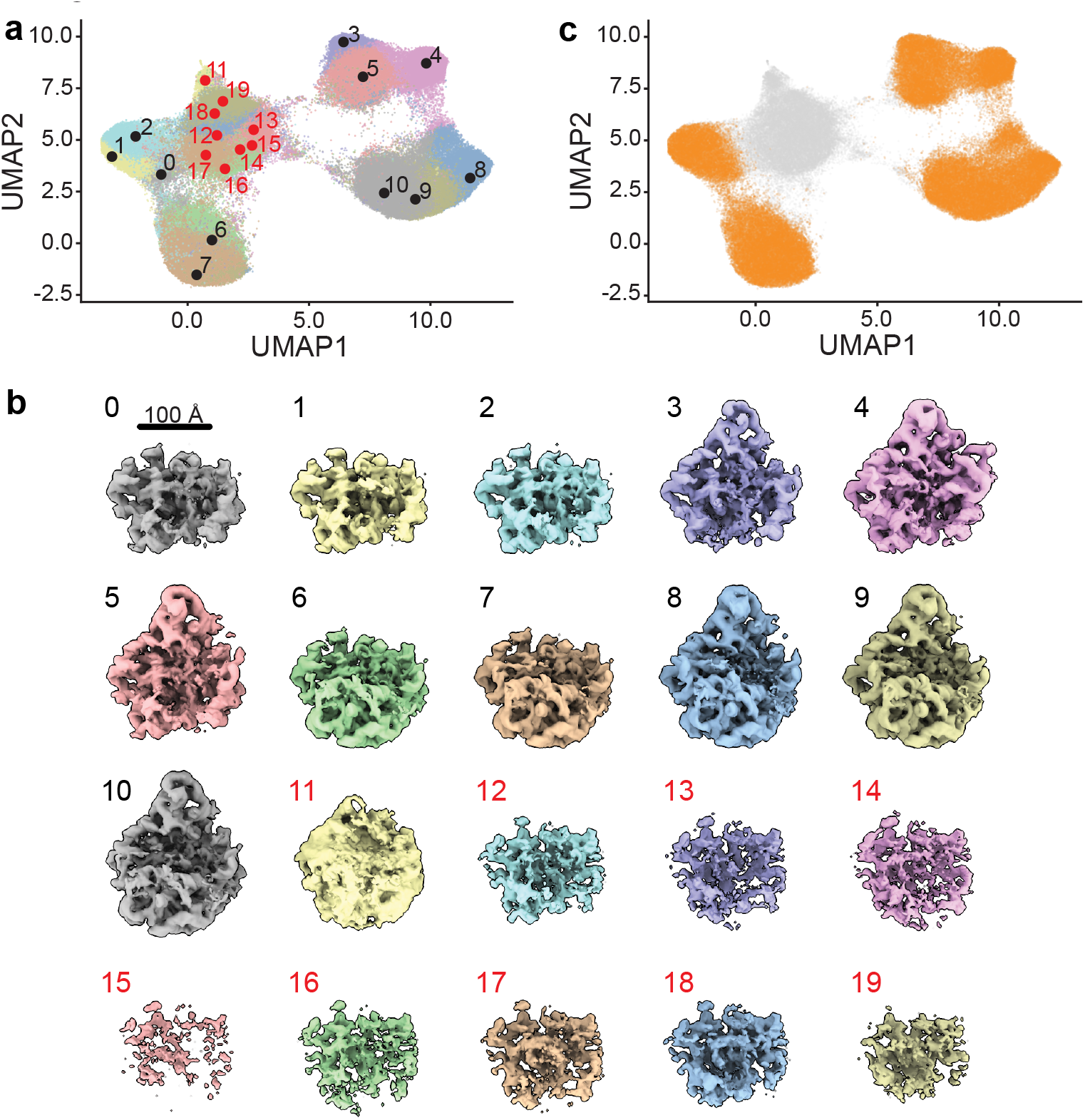
Particle filtering. **a)** UMAP visualization of latent space embeddings at epoch 49, colored by *k*-means clustering with *k*=20. Cluster centers are annotated. **b)** Volumes generated at each *k*-means cluster center. Map colors correspond to those in a. Note that volumes generated from clusters 11-19, labeled in red, are poorly resolved, consistent with the presence of poor-quality particles in the particle stack. **c)** UMAP embedding of particles selected for further training colored orange, and poor-quality particles excluded from further training are colored gray.

To facilitate this process, we provide a Jupyter notebook, `cryoDRGN_filtering.ipynb`, that allows users to filter particles interactively or using automated selections based on features (*e.g.* clusters or z-scores) of the particle embeddings (**Box 2**). The choice of particle filtering method is dataset-specific; datasets with highly featured latent representations may be more amenable to filtering by clustering or interactive selection, whereas datasets with less featured latent representations may require filtering based on z-score outliers. Particle images selected by any of these approaches can be visualized within the notebook, enabling users to examine these particles manually and adjust their selections (**Extended Data Figure 3**). We also provide tools to export the selected particles to cryoSPARC or RELION to further assess particle subsets by 2D-classification or traditional 3D-reconstruction. Lastly, this notebook allows users to save a `ind.pkl` file that records which particles have been retained (or excluded) in the filtering process. This file can be directly passed to `cryodrgn train_vae` for additional training.

##### Box 2: Particle Filtering

Several methods to filter particles are implemented in cryoDRGN. The optimal method for particle filtering is dataset-specific, and users are encouraged to try several methods to determine which is the best-suited for their particular data. We recommend that all particle filtering pipelines, regardless of method employed, start with visually inspecting *k*-means cluster center volumes generated automatically by `cryodrgn analyze`, and cross-referencing these to the UMAP plots of the *k*-means clusters in the `cryoDRGN_filtering.ipynb` notebook. Volumes from within clusters containing particle picking artifacts often appear noisy or have particularly weak density. When users have determined which regions of latent space appear to represent such artifacts, they can proceed to use any of the following methods in the Jupyter notebook to exclude particles belonging to these regions:

###### Filtering by clustering

In cases where users are able to clearly identify undesired clusters using the *k*-means cluster center volumes, they can directly select these clusters to be filtered out within the `cryoDRGN_filtering.ipynb` notebook. Gaussian mixture model (GMM) clustering can also be used, as described in this protocol.

###### Filtering by interactive selection

If there is a clearly-defined region of undesired particles within latent space, users may find it easiest to use the interactive widget in the `cryoDRGN_filtering.ipynb` notebook to manually select this region of latent space via a lasso tool and filter out all particles contained within it.

###### Filtering on magnitude of the latent embeddings

In some datasets, “junk” particles can be easily distinguished by outlying latent embedding values. This may be particularly valuable for datasets with less featured latent spaces, where the regions corresponding to particle picking artifacts are less amenable to separation by clustering or interactive selection. With the filtering notebook, users can compute the magnitude of the latent embedding and eliminate particles for which the magnitude is more than a defined number of standard deviations above the mean.

Particle filtering efficacy can be assessed by several metrics, including generating more volumes from regions of latent space enriched for retained or discarded particles and confirming the presence of good volumes and poor volumes, respectively. Users can also directly view particles in the `cryoDRGN_filtering.ipynb` notebook to see if they contain ice or edge artifacts, or other contaminants unrelated to the complex of interest. Finally, retained and discarded particles can be exported and further inspected via traditional 2D classification or 3D reconstruction in other processing software such as cryoSPARC or RELION. See **Extended Data Figure 3** for a comparison of these particle filtering methods using the EMPIAR-10076 dataset; note that all three filtering methods identify largely overlapping particle sets for this dataset.

#### Interactively exploring cryoDRGN models

Generally, analysis of a cryoDRGN model involves both visualizing the latent embeddings of particle images and generating volumes from the latent representation to understand the structural heterogeneity within the dataset (**Figure 4**). The `cryodrgn analyze` command automates these tasks by: 1) performing principal component analysis (PCA) and Uniform Manifold Approximation and Projection (UMAP; McInnes et al., 2018) on the latent embeddings to aid in visualization of high (>2) dimensional latent spaces (see **Glossary**); 2) generating representative volumes from different regions of the latent space; and 3) producing representative structural trajectories through latent space.

**Figure 4.**
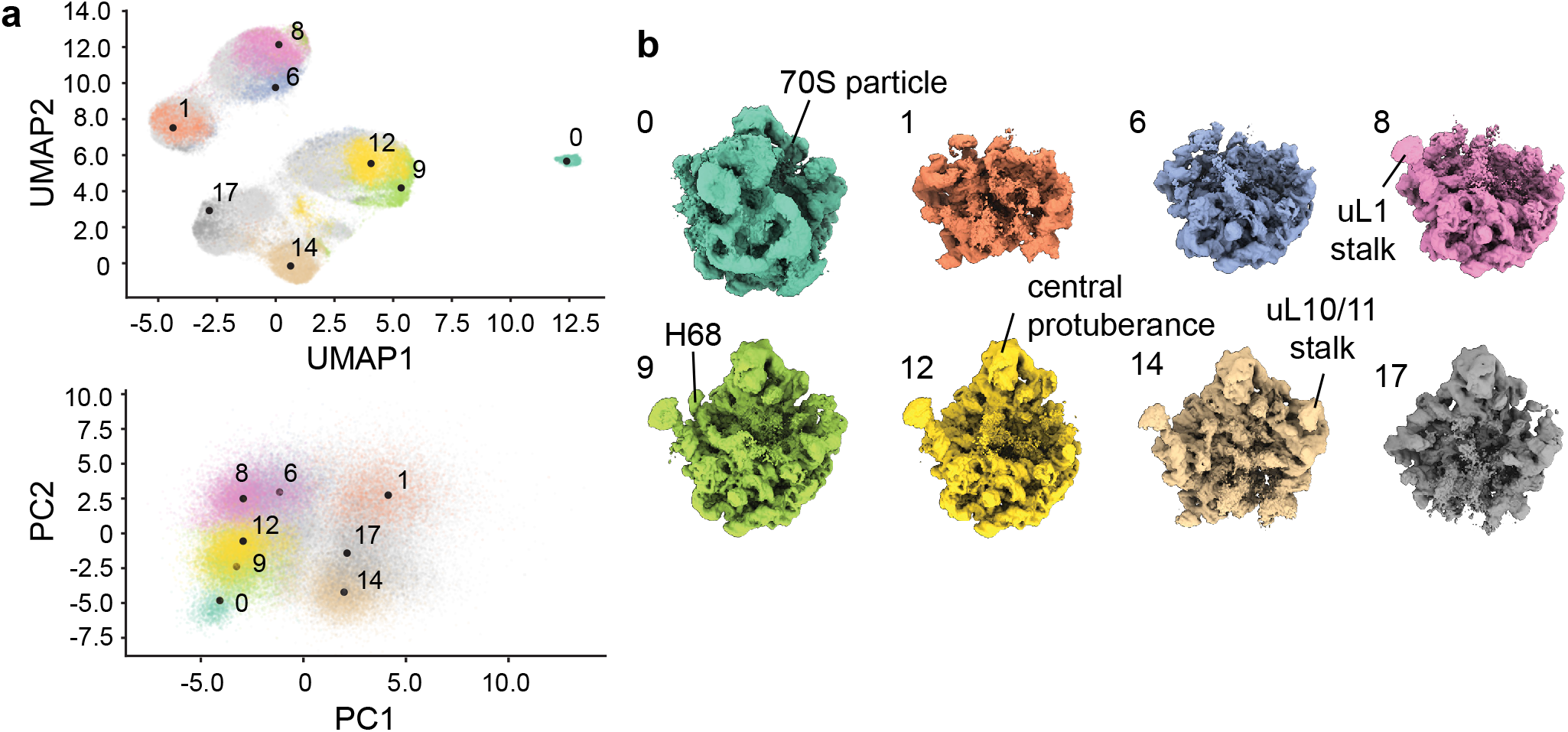
Analysis of a cryoDRGN model trained on high-resolution particle images. **a)** 8-D latent space visualized in 2-D using UMAP (*top*) or PCA (*bottom). K*-means clustering of the latent space embeddings with *k*=20 was applied, and notable clusters are colored and annotated. **b)** Representative volumes generated by the decoder network from notable cluster centers, with colors and annotation corresponding to those in a. Key structural elements of the bacterial ribosome are noted.

To visualize a representative set of volumes, `cryodrgn analyze` generates volumes after *k*-means clustering of the latent embeddings — volumes are generated at the centroid location of each *k*-means cluster. Importantly, the *k*-means algorithm is not used to identify clusters or assign particles to classes, but, instead, is used to partition the latent space into *k* regions. By default, *k*=20, and the number of sampled density maps may be modified by passing the `--ksample` argument to `cryodrgn analyze`.

To visualize representative structural trajectories, `cryodrgn analyze` generates structures along the first two principal component axes of the latent space, and trajectories along more latent dimensions can be generated using the `cryodrgn pc_traversal` tool. By default, the volumes are generated at equally spaced points between the 1^st^ and 99^th^ percentile of the data distribution projected onto each principal component. The PC trajectories can highlight major modes of the variability in the structure, however, the principal components of cryoDRGN’s latent space are not equivalent to the principal components of the volumes due to the nonlinear nature of the decoder. This contrasts with tools such as cryoSPARC’s 3DVA (Punjani and Fleet, 2021b). Additionally, it is important to note that the PC trajectories do not necessarily reflect biologically meaningful paths, and that not all volumes along these paths will originate from regions of the latent space that are well supported by data.

In addition to visualizing structures present within their dataset, users may also wish to interpolate between two or more such structures. The `cryodrgn graph_traversal` command provides a means of doing so by building a nearest-neighbor graph between all particles’ latent embeddings, finding the shortest path on the graph between specified particles, and generating volumes along the visited nodes. Unlike standard approaches that naively morph between end-point volumes or interpolate along a principal component of variability independent of underlying data support, all of the structures produced by this traversal approach are supported by data from the input particle stack, and thus may represent a more probable structural trajectory.

The `cryodrgn analyze` command also produces a Jupyter notebook, `cryoDRGN_viz.ipynb`, for interactive analysis. This notebook can be used to analyze the latent space in greater detail, to generate volumes at selected points of interest in the latent space, and to export subsets of particles for traditional 3D reconstruction and model building using other tools.

#### Interpreting structural ensembles

While `cryodrgn analyze` provides an initial characterization of the heterogeneity present within the dataset, a more systematic interrogation of the learned structures allows one to fully explore and quantify the structural states present. There are several possible approaches to perform this more systematic sampling and structural characterization, each leveraging cryoDRGN’s ability to generate large numbers of data-supported density maps. We have found supervised *“subunit occupancy analysis”* (Davis et al., 2016; Davis and Williamson, 2017) to be particularly informative for compositionally heterogeneous datasets. With this method, users designate structural elements of interest (*e.g.* RNA helices, protein complex subunits, polypeptide chains, or elements of protein secondary structure) within an aligned atomic model, and then quantify the presence or absence of each element across the structural ensemble produced by cryoDRGN. Hierarchical clustering of the resulting structural element occupancy matrix creates a highly interpretable visualization of the overall compositional heterogeneity across the structural ensemble, and reveals patterns in how these subunits are occupied, including positive and negative cooperative occupancy between individual structural elements (**Figure 5**).

**Figure 5.**
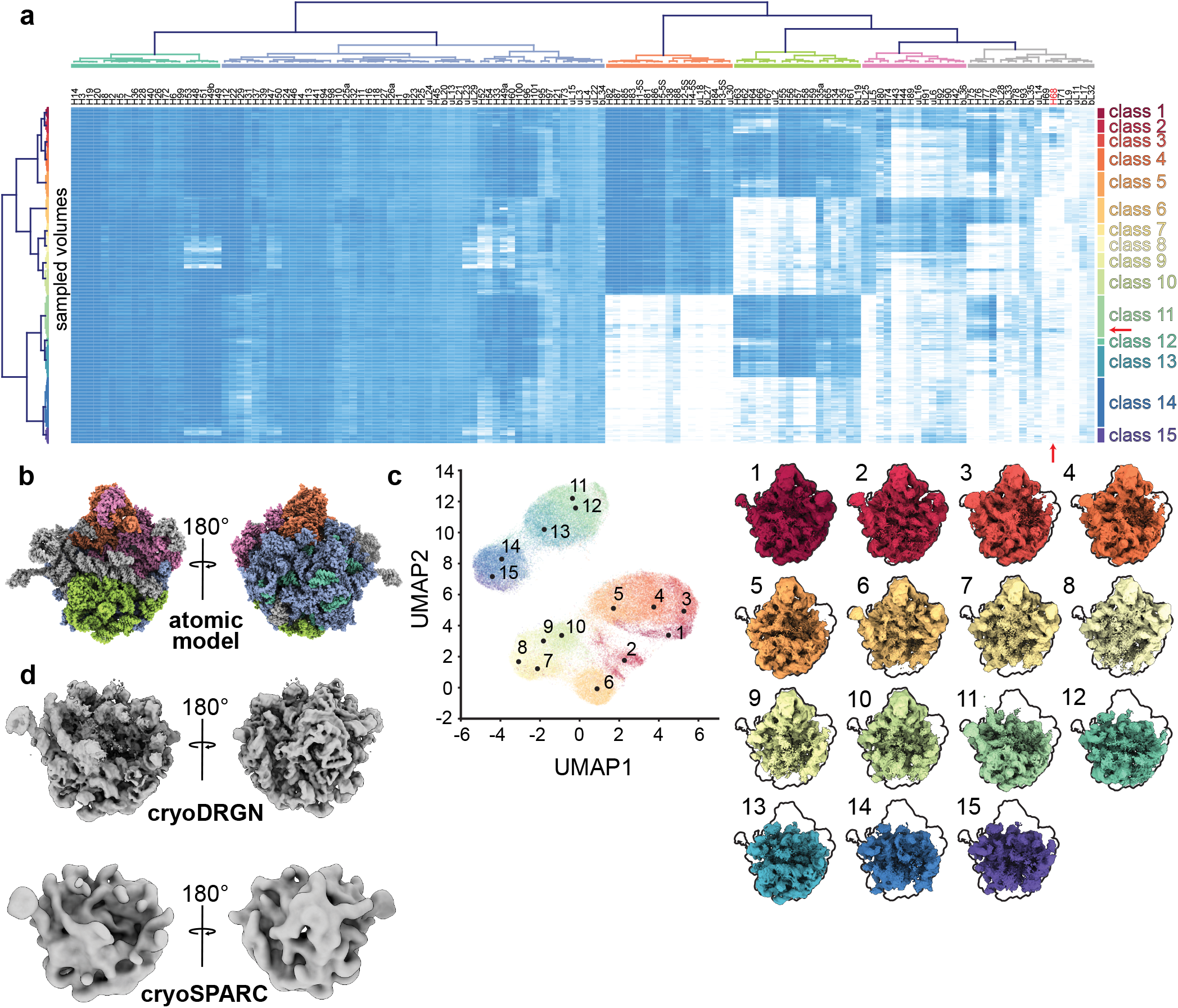
Atomic model-based analysis of cryoDRGN-generated structural ensemble. **a)** *“Occupancy analysis”* heatmap illustrating low (white) and high (blue) occupancy proteins or rRNA helices (columns) in various cryoDRGN generated density maps (rows). Using a fixed threshold linkage distance, dendrograms are colored according to structural blocks (*top*) and volume classes (*right).* A red arrow indicates the position of H68 in the heatmap. **b)** Atomic model (PDB: 4YBB) colored by structural blocks as defined in a. **c)** Centroid volumes of the occupancy analysis classes, generated at the closest on-data point to the median position in latent space for each class. Volumes are outlined for comparison to the mature 50S ribosomal subunit (class 1). **d)** C4 class example volume generated by cryoDRGN (*top*) compared to the cryoSPARC homogeneous refinement (*bottom*) using the 1,149 particles identified as class C4 through occupancy analysis. Particle group rows within class 11, and H68 column are noted with red arrows.

We note that this subunit occupancy analysis is limited by the underlying assumption that an appropriate atomic model exists and that subunits occupy their native conformation within a complex. When complexes exhibit conformational heterogeneity, it may be necessary to fit ensembles of atomic models using tools such as Molecular Dynamics-based Flexible Fitting (Trabuco et al., 2009).

In sum, this protocol provides users with a guided framework to analyze and interpret a richly heterogeneous dataset, and we expect that the approaches and tools described herein will be broadly applicable to the analysis of other datasets.

##### Glossary

###### Network architecture

the arrangement of hidden layers and nodes in each neural network. For example, a 256×3 network architecture has 3 hidden layers each containing 256 nodes, whereas the 1024×3 architecture has 3 hidden layers each containing 1024 nodes. These descriptions do not include the nodes in the input or output layers, as these are determined by the image size and by the dimensionality of the latent space.

###### Encoder network

the neural network that encodes each particle image in a low-dimensional latent space. By default, we use an 8-dimensional latent space, though users can specify higher or lower dimensions.

###### Decoder network

the neural network that generates a 3-D density map, given a latent embedding.

###### Epoch

the passage of an entire particle stack through the encoder and decoder networks. The networks are iteratively trained through multiple epochs.

###### Minibatch

Particles are passed through the encoder and decoder networks in groups called mini-batches of 8 images by default; changing the mini-batch size affects memory utilization, training dynamics, and training speed.

###### PCA

a *linear* dimensionality technique used in this protocol to visualize the latent space. Axes produced by PCA are orthogonal and ordered by maximum variance along each axis, and we typically inspect the first 2-4 axes. In practice, we find that PCA is useful for identifying outliers in the latent embedding distribution and summarizing major modes of heterogeneity, however we find that useful local structure in the distribution is often lost due to the linear projection.

###### UMAP

a *non-linear* dimensionality reduction technique used in this protocol to embed the latent space into an easily visualized 2-D space. UMAP tends to highlight local neighborhood structure at the expense of preserving global structure. As a result, distance metrics in UMAP-space such as inter-cluster distance are not generally meaningful. We find that UMAP embeddings are useful in segmenting structurally disparate groups of particles and that high particle densities within a UMAP cluster meaningfully represent dense particle neighborhoods in latent space.

###### Z-score

the number of standard deviations above the mean. Used during particle filtering to identify particles with a z-score > 2 (by default), meaning a latent embedding whose magnitude is 2 standard deviations above the mean magnitude across all particles.

###### On-data

volumes generated by the described cryoDRGN analysis scripts are always generated from a position in the latent space directly corresponding to the latent embedding of some particle within the input stack – *i.e.*“on-data” Specifically, we generate a volume from the closest on-data point to a given query in latent space.

###### UMAP local maxima method

our approach to identify a set of latent coordinates representing diverse particles in areas of latent space that are well-supported by data. This method aims to automatically reproduce how a user might interactively select a subset of dense clusters from a UMAP embedding. Briefly, latent values for all particles from the final epoch of training are embedded in 2-D UMAP space. This space is then binned with 30 bins per axis and the resulting 2-D histogram is smoothed with a Gaussian of width = 1 bin. All local maxima are identified, then greedily pruned such that the lower amplitude maximum of two local maxima within a defined radius of each other is removed. A final filtering step returns the 10 largest local maxima. Particles within a 3×3 grid of bins centered on each local maximum are labeled as corresponding to local maxima A-J, and their on-data median latent coordinate is returned for volume generation. Note that maxima are labeled A-J in order of decreasing particle count.

## MATERIALS

### Equipment

The minimal compute requirements for this protocol are as follows:

- **Linux workstation or cluster:** equipped with at least one NVIDIA GPU (Pascal, Turing, Volta, and Ampere architectures have been tested), 128 GB RAM, and 250 GB disk space for all raw data and outputs.

Performance will vary based on system configuration. For compute-expensive steps of the protocol, we provide approximate timings using a system equipped as follows:

- **CPU:** Dual Xeon Gold 6242R processors
- **GPU:** NVIDIA 3090 RTX (single GPU)
- **RAM:**512 GB

### Requisite software

- **CryoDRGN installation:** Updated installation and usage instructions are maintained at https://github.com/zhonge/cryodrgn. Installation instructions for cryoDRGN v0.3.5 which is used in this paper can be found in Supplementary Protocol 1.
- **UCSF ChimeraX**: Installation instructions are at https://www.cgl.ucsf.edu/chimera/download.html
- **RELION v3.1.1**: Installation instructions are at https://relion.readthedocs.io/en/latest/Installation.html
- **cryoSPARC** (*optional*): Version 2.4 from https://cryosparc.com/download was used in this protocol.
- **Occupancy analysis:** Installation instructions, segmented .pdb files, Python and shell scripts, and Jupyter notebook for analysis available at https://github.com/lkinman/occupancy-analysis. Version 0.1.2 was used in this protocol.

### Requisite datasets

- **EMPIAR-10076 particle stack:** Download from EMPIAR web interface at: https://www.ebi.ac.uk/empiar/EMPIAR-10076/ or via the command:

~~~
rsync -avzP empiar.pdbj.org::empiar/archive/10076 ./
~~~
- **EMPIAR-10076 reconstruction metadata:** Download 00_inputs from https://doi.org/10.5281/zeno-do.5164127, or follow the cryoSPARC reconstruction guide in Supplementary Protocol 2.
- **Precomputed results** (*optional*): All results from this protocol can be downloaded via a web browser from https://doi.org/10.5281/zenodo.5164127. Files can alternatively be downloaded from the command line:

~~~
pip install zenodo_get
zenodo_get --md5 -w urls.txt 5164127
wget -c -i urls.txt
md5sum -c md5sums.txt
conda install zstd
for i in *.tar.zst; do tar --use-compress-program=unzstd -xvf ${i}; done
~~~

## PROCEDURE

### Prepare cryoDRGN inputs [40 minutes]

1. Connect to the workstation containing the cryoDRGN installation and all downloaded files. We will assume that the base directory contains the downloaded EMPIAR-10076 particle stack at `./10076/data/L17Combine_weight_local.mrc` and the downloaded reconstruction and reconstruction metadata at `./00_inputs/cryosparc_P71_J21_004_volume_map.mrc` and `./00_inputs/ cryosparc_P4_J33_004_particles.cs`, respectively. The majority of the commands in this protocol will be run within this base directory. Ensure the cryoDRGN conda environment is activated, which is required for all `cryodrgn` commands. Enter the following at the terminal.

~~~
cd /path/to/base/directory
conda activate cryodrgn
cryodrgn --help
~~~ The last line provides a list of possible cryoDRGN commands. To learn more about a particular cryoDRGN command, simply enter `cryodrgn [command] --help`.
2. Convert poses^1^ from the downloaded cryoSPARC refinement to cryoDRGN format as `poses. pkl`, specifying the refinement box size of D=320px. Rename the `particles.cs` file appropriately if you ran your own cryoSPARC refinement following Supplementary Protocol 2.

~~~
cryodrgn parse_pose_csparc --help
cryodrgn parse_pose_csparc 00_inputs/cryosparc_P4_J33_004_particles. cs -D 320 -o poses.pkl
~~~ CryoDRGN will report pose information for the first particle as well as the number of particles parsed (131,889).
3. Convert CTF parameters^1^ from the downloaded cryoSPARC refinement to cryoDRGN format as `ctf.pkl`.

~~~
cryodrgn parse_ctf_csparc 00_inputs/cryosparc_P4_J33_004_particles. cs -o ctf.pkl
~~~ CryoDRGN will report relevant imaging parameters including image size and pixel size.
4. Downsample the dataset to D=128px and D=256px, which we will use in the first and second training rounds, respectively. Here we split the particle stacks into 50,000 particle sub-stacks with `--chunk` to decrease the memory footprint. These chunked .mrcs files are identified by an auto-generated `particles.[px].txt` file.

~~~
cryodrgn downsample 10076/data/L17Combine_weight_local.mrc -D 128 -o particles.128.mrcs --chunk 50000
cryodrgn downsample 10076/data/L17Combine_weight_local.mrc -D 256 -o particles.256.mrcs --chunk 50000
~~~
5. Verify that all data was parsed correctly by back-projecting the first 10,000 particles and comparing the resulting map (`backproject.128.mrc`) with the refined consensus map `cryosparc_P71_J21_004_volume_map.mrc` using ChimeraX. Overall, the back-projected map should match the refined map well, albeit at lower resolution and with more noise (**Extended Data Figure 1**).

~~~
cryodrgn backproject_voxel particles.128.txt --uninvert-data --poses poses.pkl --ctf ctf.pkl -o backproject.128.mrc
~~~

**Extended Data Figure 1.**
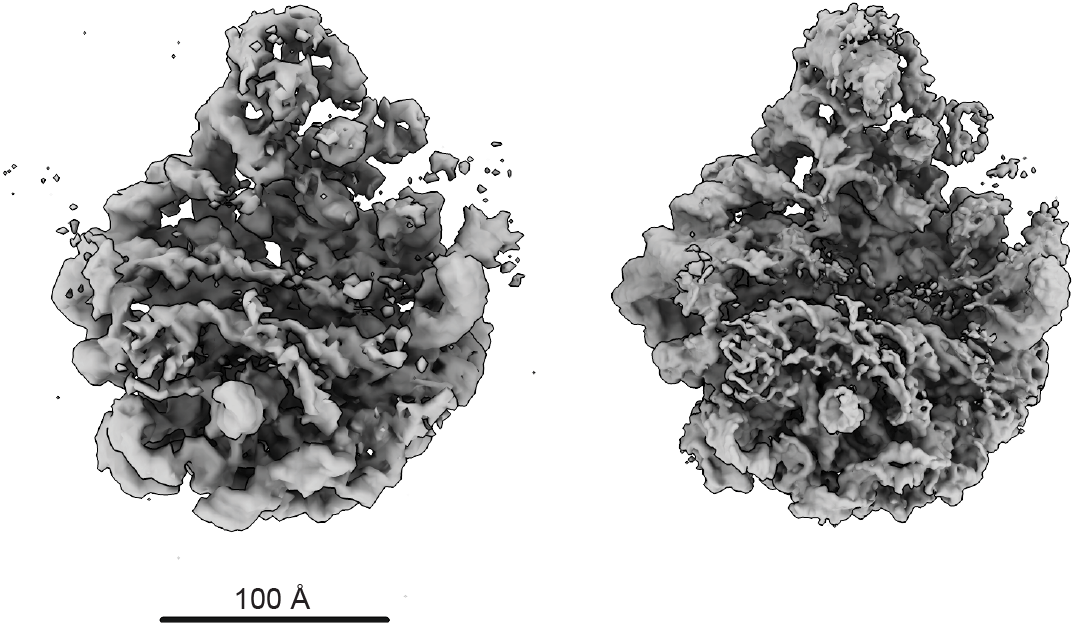
Assessing cryoDRGN input parsing. Comparison of 10,000 back projected cryoDRGN-parsed particles at D=128px (*left*) with the unsharpened map from cryoSPARC’s homogeneous refinement (*right*).

### Train cryoDRGN networks [5 hours]

6. Run the `cryodrgn train_vae` command to begin training. To expedite this initial training run, we use a small neural network architecture (256×3) on the 128px downsampled particles. A description of all available training parameters can be displayed with cryodrgn `train_vae --help`. If memory utilization is limiting on the GPU, decreasing the `--batch-size` parameter (see **Glossary**) from its default value of 8 may be helpful. However, training dynamics will also be affected and additional training epochs may be required. Note also that the AMP acceleration used here is most effective for large network architectures and image sizes that are divisible by 8.

~~~
cryodrgn train_vae particles.128.txt --ctf ctf.pkl --poses poses.pkl --zdim 8 -n 50 --batch-size 8 --amp --uninvert-data -o 01_128_8D_256 > 01_128_8D_256.log &
~~~ **Pausepoint:** the training command will take several hours to run, depending on your computational hardware. You can follow training progress with `tail -f 01_128_8D_256.log`. If your job is interrupted or you otherwise want to restart or extend training, you can resume from any epoch by adding `--load 01_128_8D_256/weights.[EPOCH#].pkl` to the above `train_vae` command. Alternatively, you can specify `--load_latest` to avoid providing the specific path to the `weights.pkl` file for the most recent epoch.
7. After 50 epochs of training have completed, we check if the network has converged such that additional training would not be beneficial.

~~~
python /path/to/cryodrgn/utils/analyze_convergence.py --help
python /path/to/cryodrgn/utils/analyze_convergence.py 01_128_8D_256 49 --flip
~~~ **Critical:** the `--flip` flag used here changes the handedness of the map and should be set accordingly in all subsequent steps that generate volumes. If you generated your own consensus reconstruction, visualize your map to determine if this flag is required. All outputs are saved to `01_128_8D_256/convergence.49`, including plots of each heuristic convergence metric (**Figure 2; Extended Data Figure 2**). A description of the purpose, implementation, and interpretation of all convergence heuristics is included in **Box 1**. For each heuristic, a plateau is consistent with convergence and, although individual metrics can be noisy and vary in a dataset-dependent manner, we define convergence as the epoch upon which most of these metrics have plateaued. For this training job, we examine the plots in `01_128_8D_256/ convergence.49/plots` and observe convergence between epochs 29 and 39. These plots also give insight into training dynamics and the distribution of latent variable embeddings. For example, the latent embedding of this dataset is highly featured with clearly visible clusters in the UMAP embedding plots. This will not be the case for all datasets, and many datasets exhibit a less featured latent space, yet display significant heterogeneity when volumes are visualized from distal points in the latent space. At this point, users may find it useful to examine the volumes in `convergence.49/vols.[EPOCH#]` directly for evidence of “overtraining” pathologies such as increased noise or streaking artifacts. If overtraining artifacts are observed, users should examine volumes from earlier training epochs, as they may lack such pathologies.
8. In the event that training has not converged, users should return to step 6 (`cryodrgn train_vae`), increase the number of epochs to 100 (i.e. `-n 100`), and load the epoch 49 weights with `--load 01_128_8D_256/weights.49.pkl` to resume training. Users should then reassess convergence as above before proceeding with particle filtering and model analysis. In general, the number of epochs should be chosen with the dataset size and suitable training time in mind.

**Extended Data Figure 2.**
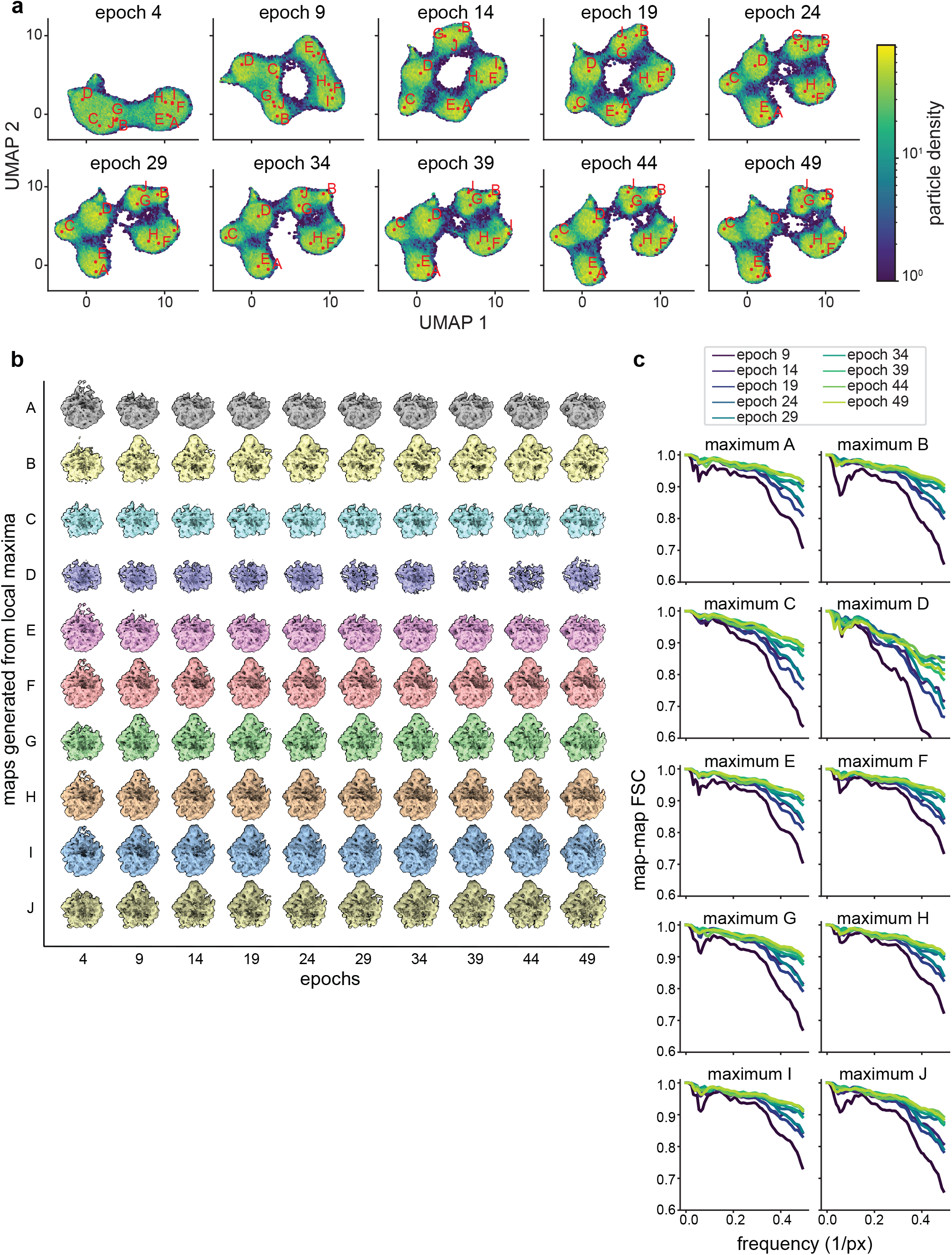
Assessing convergence of representative cryoDRGN density maps during network training. **a)** Particle sets of interest A-J identified in epoch 49 by the *“UMAP local maxima method”* (Glossary) are mapped to prior epochs’ UMAP embeddings. The on-data median latent value of each particle set is embedded into UMAP space and annotated for each epoch. Note that each annotated point maps to the same high occupancy region of UMAP space following convergence. **b)** Corresponding volumes generated from each on-data median latent value at five epoch intervals as shown in panel a. Note that the volumes’ gross morphology stabilizes by epoch 14-19, though some additional details in maxima I and J require 2429 epochs of training. **c)** FSC plots correlating each local maximum volume at epochj and at epoch_j-5_.

### Filter particles [30 minutes]

9. Run the `cryodrgn analyze` command to perform an automated analysis of the trained cryoDRGN model at epoch 49:

~~~
cryodrgn analyze 01_128_8D_256 49 --flip --Apix 3.275
~~~ This command runs PCA and UMAP on the embedded latent space, generates volumes at 20 *k*-means cluster centers, and creates interactive Jupyter notebooks for further visualization and analysis. Note that the user should provide the pixel size in angstroms after accounting for downsampling (`--Apix`). If the user wishes to change the number of *k*-means cluster center volumes generated, this can be accomplished using the `--ksample` flag. The `--flip` flag is set to invert volume chirality, as described above. Note that all volumes generated in this protocol are generated from “on-data” positions in latent space (see **Glossary**).
10. A new directory named analyze.49 was created within `01_12_8_8D_256` by `cryodrgn analyze`. The directory contains subdirectories `pc1, pc2`, and `kmeans20`, along with plots of the latent space (`umap.png` and `z_pca.png`) and the Jupyter notebook files `cryoDRGN_filtering.ipynb` and `cryoDRGN_viz.ipynb`. View the contents of this folder to verify that the analysis script ran correctly.
11. Before proceeding to high-resolution training, we will eliminate contaminating particles including edge artifacts and ice. Here, we demonstrate one possible filtering approach using a combination of *k*-means and Gaussian mixture model (GMM) clustering of latent space. Additional approaches involving manual selection or filtering on outlying latent values are implemented within the notebook, and may be more appropriate for other datasets (**Box 2; Extended Data Figure 3**). Open the `cryoDRGN_filtering.ipynb` notebook within Jupyter Lab. See **Supplementary Protocol 3** for notes on how to access this Jupyter notebook remotely if you are running this protocol on a compute cluster.
12. Check that `EPOCH=49` in the third cell is set and run all cells up to the *“Filter by cluster”* header.
13. Examine the volumes generated by *k*=20 *k*-means clustering using ChimeraX by opening all volumes in the `01_128_8D_256/analyze.49/kmeans20` directory. Comparing the centroid locations of these clusters in UMAP space, as visualized within the notebook, to their corresponding volumes suggests that for this dataset the central teardrop-shaped UMAP cluster containing *k*-mean centers 11-19 contains primarily poor-quality particles (**Figure 3A-B**). This conclusion is based on the appearance of the volumes, with the volumes that fall within the “junk cluster” showing significant noise (11) or much weaker density than the other volumes (12-19) at a uniform isosurface level. Note that due to random initialization each time cryoDRGN and UMAP are run, users may find this cluster differently shaped, comprised of different *k*-means classes, and in a different area of their UMAP space. We note that the number and latent distribution of junk particles is highly dataset-specific, and identification of which particles should be filtered often requires detailed inspection of the volumes by the users, and is typically verified by subsequent inspection of representative particles and corresponding cryoDRGN volumes, and by traditional 2-D classification or 3-D reconstructions using these particles stacks.
14. To exclude these particles from further analysis, move to the GMM clustering section of the notebook. Run GMM clustering with `G=5` or `G=6` such that the plot of UMAP space colored by GMM cluster shows a clean separation of the junk cluster (here, the central teardrop-shaped cluster) from the rest. Due to GMM’s random initialization, this may take several iterations and may separate as one or two GMM clusters (**Figure 3C**). Alternatively, this cluster may be specified by the corresponding *k*-means clusters (here, 11-19), or by selection with a lasso tool in the interactive selection section of the filtering notebook (**Box 2**).
15. In the subsequent cells, define `ind_selected` as an array of GMM cluster labels that exclude the junk particle cluster (for our initialization, cluster 3), such as `ind_selected=[0,1,2,4,5]`.
16. Under the *“Save selection”* header, run all cells to save the filtered particle indices as a `.pkl` file that can be fed directly to `cryodrgn train_vae`. We expect to have included ~100,000 particles and excluded ~30,000. Alternatively, download the filtered particle indices from precomputed-`01_128_8D_256.tar.zst` as described in **Materials**.
17. Optionally, a .star file corresponding to these filtered indices can be created at the command line for import into traditional reconstruction software. For example, export both good and bad particles to assess particle filtering by 2D-classification in cryoSPARC (**Extended Data Figure 3**) using the `write_star.py` script found within the cloned cryoDRGN software directory:

~~~
mv 10076/data/L17Combine_weight_local.mrc 10076/data/L17Combine_weight_local.mrcs
cryodrgn write_starfile 10076/data/L17Combine_weight_local.mrcs ctf. pkl --poses poses.pkl --ind 01_128_8D_256/ind_keep.96478_particles. pkl -o ind_keep.star
cryodrgn write_starfile 10076/data/L17Combine_weight_local.mrcs ctf. pkl --poses poses.pkl --ind 01_128_8D_256/ind_bad.35421_particles. pkl -o ind_bad.star
~~~

**Extended Data Figure 3.**
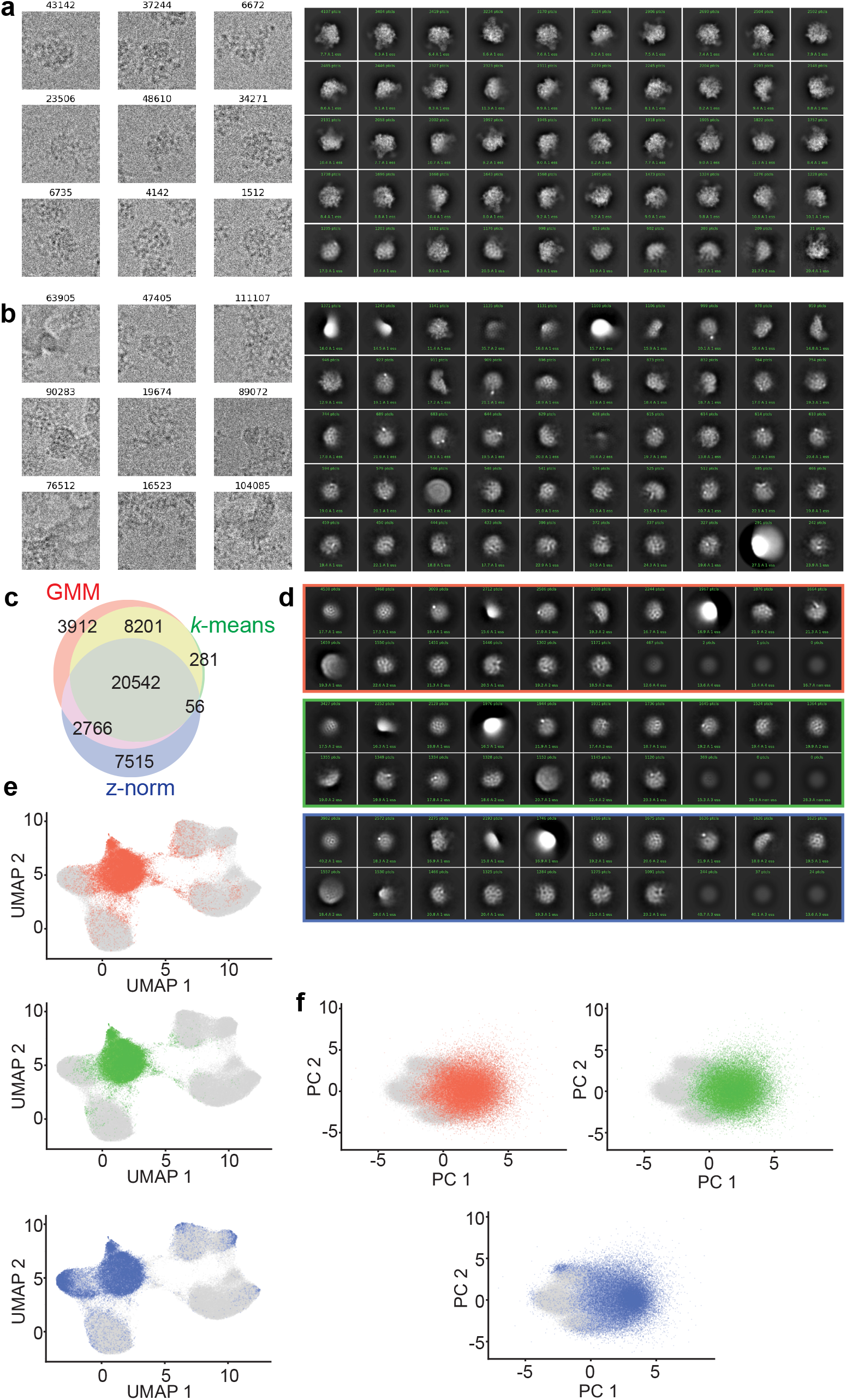
Visualizing particle filtering. **a)** Representative particles filtered by `ind_keep.star`, selected for further training, and corresponding 2D-classification using default cryoSPARC parameters. **b)** Representative particles filtered by `ind_bad.star`, excluded from further training, and corresponding 2D-classification using default cryoSPARC parameters. **c)** Three-way Venn diagram of “junk” particles identified by one of the following methods: two classes from *k*=6 Gaussian mixture model latent-space classification (red, 35,421 particles); ten classes from *k*=20 *k*-means latent-space classification (green, 29,080 particles); or latent encoding magnitude (z-norm) exceeding 0.5 standard deviations larger than the mean (blue, 30,879 particles). **d)** Corresponding CryoSPARC 2D-classification results using “junk” particles identified through the GMM (top), *k*-means (middle), or z-norm (bottom) filtering approaches. **e)** UMAP embedding or **f)** PCA projections highlighting location of junk particles identified by GMM (red), *k*-means (green), or z-norm (blue) methods.

### Train high-resolution cryoDRGN networks [1 day]

18. Train a new model using the particles selected above at higher resolution (D=256px) and using a larger network architecture (1024×3). Training this model is the most computationally expensive step in this protocol, and using `--amp` decreases training times. If your workstation is equipped with less than 128 GB RAM, you may encounter out-of-memory errors. These errors can be circumvented by the addition of `--lazy` to the following command, allowing on-the-fly image loading from disk at a significant cost of performance.

~~~
cryodrgn train_vae particles.256.txt --amp --ctf ctf.pkl --poses poses.pkl --zdim 8 -n 50 --uninvert-data --enc-dim 1024 --enc-layers 3 --dec-dim 1024 --dec-layers 3 -o 02_256_8D_1024 --ind 01_128_8D_256/ ind_keep.96478_particles.pkl > 02_256_8D_1024.log &
~~~ **Pausepoint:** as in step 6, this training step will require many hours to complete.
19. Verify that the new model has converged by running:

~~~
python /path/to/cryodrgn/utils/analyze_convergence.py 02_256_8D_1024 49 --flip
~~~ We find that the network has satisfactorily converged by the end of training according to the criteria described in step 7 above (**Extended Data Figures 4,5**).

**Extended Data Figure 4.**
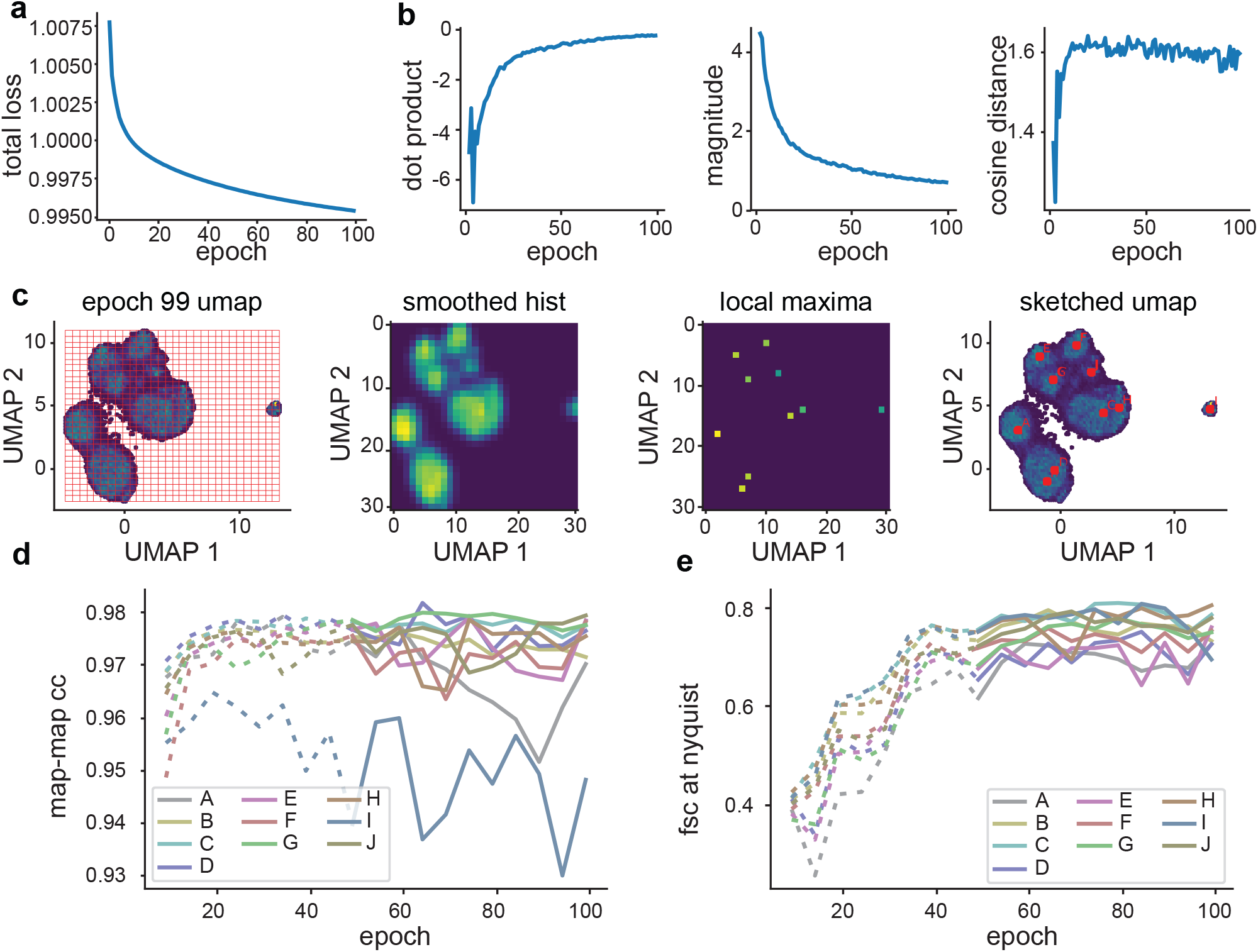
Training and assessing convergence of high-resolution training. **a)** Representative plot of average total loss at each epoch. **b)** Median per-particle movement through latent space, characterized by vectors connecting each particle’s latent embedding in successive epochs. Resulting vector dot products (*left*), magnitude (*center*) and cosine distance (*right*) are shown. **c)** Identification of representative latent embeddings via the *“UMAP local maxima method”* (Glossary). The UMAP embedding of epoch 99 is binned into a 2-D histogram, smoothed, annotated with local maxima, and overlaid with the maxima. The on-data median UMAP location of each maximum and its neighboring 8 bins is shown. Label order corresponds to decreasing particle count in each local maximum. **d)** Map-map correlation and **e)** FSC at Nyquist limit calculated between representative volumes generated as defined in c at five epoch intervals. Epochs for which the encoder network has not converged are noted with dotted lines.

### Interactively explore cryoDRGN models [30 minutes]

20. Run `cryodrgn analyze` at the desired epoch. Here, we will analyze epoch 49 based on the convergence criteria described above.

~~~
cryodrgn analyze 02_256_8D_1024 49 --Apix 1.6375 --flip
~~~
21. Launch Jupyter Lab to explore the `cryoDRGN_viz.ipynb` file located in `02_256_8D_1024/ analyze.49`. Run the cells sequentially, verifying that epoch number 49 is entered in the third cell. This notebook produces a series of figures, some of which allow for interactive visualization of the distribution of poses, CTF parameters, or defocus values in latent space. Comparing these distributions allows us to check if the latent variable is capturing variation in non-structural heterogeneity, such as imaging parameters or viewing direction. Other figures include global pose distribution within the dataset and the latent space colored by *k*-means clusters with center points of clusters annotated (**Extended Data Figure 6**).
22. Visualize the *k*-means cluster center volumes generated by `cryodrgn analyze` in ChimeraX by opening all volumes in the `02_256_8D_1024/analyze.49/kmeans20` folder. Compare these volumes to the annotated *k*-means cluster centers in the latent space graphs (`umap.png` and `umap_hex.png` in the same directory). The latent space for this dataset is highly structured, with clusters visible by UMAP that correspond to assembly states of the ribosomal 50S subunit originally identified by Davis et al. Selected volumes and their corresponding particles are shown in **Figure 4**.
23. To help understand the major modes of motion within the dataset, visualize the volumes generated along the first two principal components in ChimeraX (**Supplementary Movies 1 and 2**). These volumes are located in the `02_256_8D_1024/analyze.49/pc1` and `02_256_8D_1024/analyze.49/pc2` subdirectories, respectively. Note that the first principal component encodes variable density in the base of the ribosome, whereas the second principal component encodes variable density in the central protuberance.

### Interrogate structural ensembles using an atomic model [2 hours]

Now that we have a sense of the types of variability present in this dataset, we seek to more systematically sample and analyze this structural heterogeneity. We use a supervised *“subunit occupancy analysis”,* as we identified extensive compositional heterogeneity in the observed k-means cluster center volumes. Here we will generate 500 volumes using *k*-means clustering and interpret their structural heterogeneity using an aligned atomic model. These 500 volumes can be generated directly by re-running the `cryodrgn analyze` command with the optional argument `--ksample 500`, or through the `cryoDRGN_viz.ipynb` interactive Jupyter notebook as described in steps 24-25 below.

24. To focus our subunit occupancy analysis on assembling large subunit particles, we first filter the small number of contaminating 70S particles that appear as outliers in the latent space (**Figure 4**). The criteria defining these particles may change from run-to-run; here, we distinguish these particles as those with UMAP1 > 10. Create a new cell in the `cryoDRGN_viz.ipynb` notebook and enter the following to perform *k*-means clustering with 500 cluster centers on the remaining particles. Users may adjust the definition of the “sub” dataframe in the first line to reflect their own criteria to exclude the 70S particles.

~~~
sub = df[df[UMAP1’] < 10]
sub_z = z[sub.index]
K = 500
kmeans_labels, centers = analysis.cluster_kmeans(sub_z, K)
centers, centers_ind = analysis.get_nearest_point (sub_z, centers)
centers_ind_df = sub.index[centers_ind]
sub.loc[:, ‘Kmeans500’] = kmeans_labels
sub.to_csv(‘kmeans500 df.csv’)
utils.save_pkl(centers_ind_df, ‘kmeans500_labels.pkl’)
~~~ Note that the last three lines save the information about which particles belong to which *k*-means cluster, and which particles within each cluster represent the cluster center. We use these data below.
25. Navigate to the *“Generate volumes”* section of the same Jupyter notebook and change the `vol_ind` definition to `vol_ind=centers_ind_df`. Additionally, several cells below this, set `Apix=1.6375` and set `flip=True`. After making these changes, run the cells in this section in order, generating 500 volumes corresponding to the on-data centers of the 500 *k*-means clusters identified in the previous step (**Extended Data Figure 7**).
26. Subunit occupancy analysis requires an aligned atomic model segmented into chains indicating the structural elements of interest. The segmented PDB files for this dataset are available at https://github.com/lkinman/occupancy-analysis, in the `protocol_examples` folder (**Materials**), and **Supplementary Protocol 4** details how to generate them. Chain assignments for each residue are provided in **Supplementary Table 1**. Note that the segmented PDB models must be aligned with an example cryoDRGN map prior to use; while the PDB models included in protocol_examples folder have been pre-aligned to the consensus refinement at `00_inputs/cryosparc_P4_J33_00_4_particles.cs`, instructions on how to align these models to an arbitrary volume in ChimeraX are provided in **Supplementary Protocol 5**. For the remaining workflow, we will assume you have stored all the downloaded occupancy analysis scripts in a subdirectory of your base directory called `03_occupancy_analysis`. This subdirectory will be your new working directory for steps 27-33. We also assume that the aligned .pdb files, along with the `reconstruct_000000` folder containing all 500 volumes sampled from latent space, are stored in `/path/to/base/directory/03_occupancy_analysis/00_aligned/`.
27. Navigate to your working directory (`/path/to/base/directory/03_occupancy_analysis`). Activate the cryodrgn conda environment as before and use the provided `gen_mrcs.sh` shell script to convert the segmented and aligned PDB files into .mrc files aligned to an example cryoDRGN map. This script is a wrapper for ChimeraX, thus `chimerax` must be installed and executable (*i.e.* `which chimerax` should return the path to the ChimeraX executable).

~~~
conda activate cryodrgn
bash protocol_examples/gen_mrcs.sh
~~~ This shell script may be adapted to use for other datasets by changing the names of the PDB files, the chains within each file, and the resolution, where the resolution used should be approximately the global FSC= 0.143 resolution of the consensus reconstruction. The output of this shell script is a directory called `01_PDB_mrc` containing a separate converted .mrc file for each of the 136 chains defined in the segmented PDB files.
28. Create masks from each of the .mrc files generated in the last step using RELION and the `gen_masks.py` script. This script is a wrapper for `relion_mask_create`, thus RELION must be activated in your current environment (*i.e.* `which relion_mask_create` should return the path to the `relion_mask_create` executable on your system).

~~~
for i in 01_PDB_mrc/*.mrc; do python gen_masks.py --mrc $i --outdir 02_mask; done
~~~ This script produces a pixel size warning that can be ignored.
29. Calculate the reference-normalized occupancies of each defined subunit in each of the 500 electron density maps sampled from latent space using the provided `calc_occupancy.py` script.

~~~
python calc_occupancy.py --mapdir 00_aligned/reconstruct_000000 --maskdir 02_mask --refdir 01_PDB_mrc
~~~
30. Launch Jupyter Lab and open the provided `occupancy_analysis.ipynb` template notebook. Change the occupancies variable in the second cell to indicate the location of the `occupancies.csv` file generated in the previous step (*e.g.* `/path/to/base/directory/03_occupancy_analysis/occupancies.csv`). If using this notebook on a different dataset or with different segmented atomic models, change the `num_volumes` variable and chains dictionary as necessary. The keys for the chains dictionary should be the names of the atomic model files, and the corresponding values should describe the identity of the chains in that file, in alphabetical order of the chains.
31. After changing the necessary variables, run the cells in order through the *“Normalization”* section. Here, we implement a normalization method in which the values are re-scaled to span a range from the tenth to ninetieth percentiles of the original data. The most appropriate normalization method may vary by dataset and users will likely need to try several methods of normalizing data to determine which best facilitates visual analysis interpretation of the resulting heatmap.
32. Hierarchical clustering allows us to group volumes that exhibit similar patterns of subunit occupancy. We define density map classes by setting a threshold distance on the dendrogram of the rows. In applying such a threshold to the dataset shown here, we observe classes of particles at varying stages of assembly, as evidenced by the presence and absence of various structural features in these different classes (**Figure 5A,C**). Applying a threshold to the dendrogram of the columns identifies structural blocks consisting of rRNA or protein elements that show similar occupancy patterns. Coloring the atomic model by these structural blocks improves the interpretability of the clustermap and assists in visualizing cooperative blocks that may be present (**Figure 5A,B**). Clustering can be performed within the provided Jupyter notebook by running the *“Hierarchical clustering”* cells.
33. Run the *“Extract classes from clustering”* section of the notebook to automatically extract the volume classes and structural blocks at the thresholds you defined in step 32. The subsequent two sections of the notebook, *“Visualize volume classes in ChimeraX”* and *“Visualize structural blocks in ChimeraX”,* produce a series of .py scripts that can be opened in ChimeraX for direct visualization of the subset of kmeans-500 electron density maps within each volume class, and the atomic model colored by the structural blocks, respectively.

**Extended Data Figure 5.**
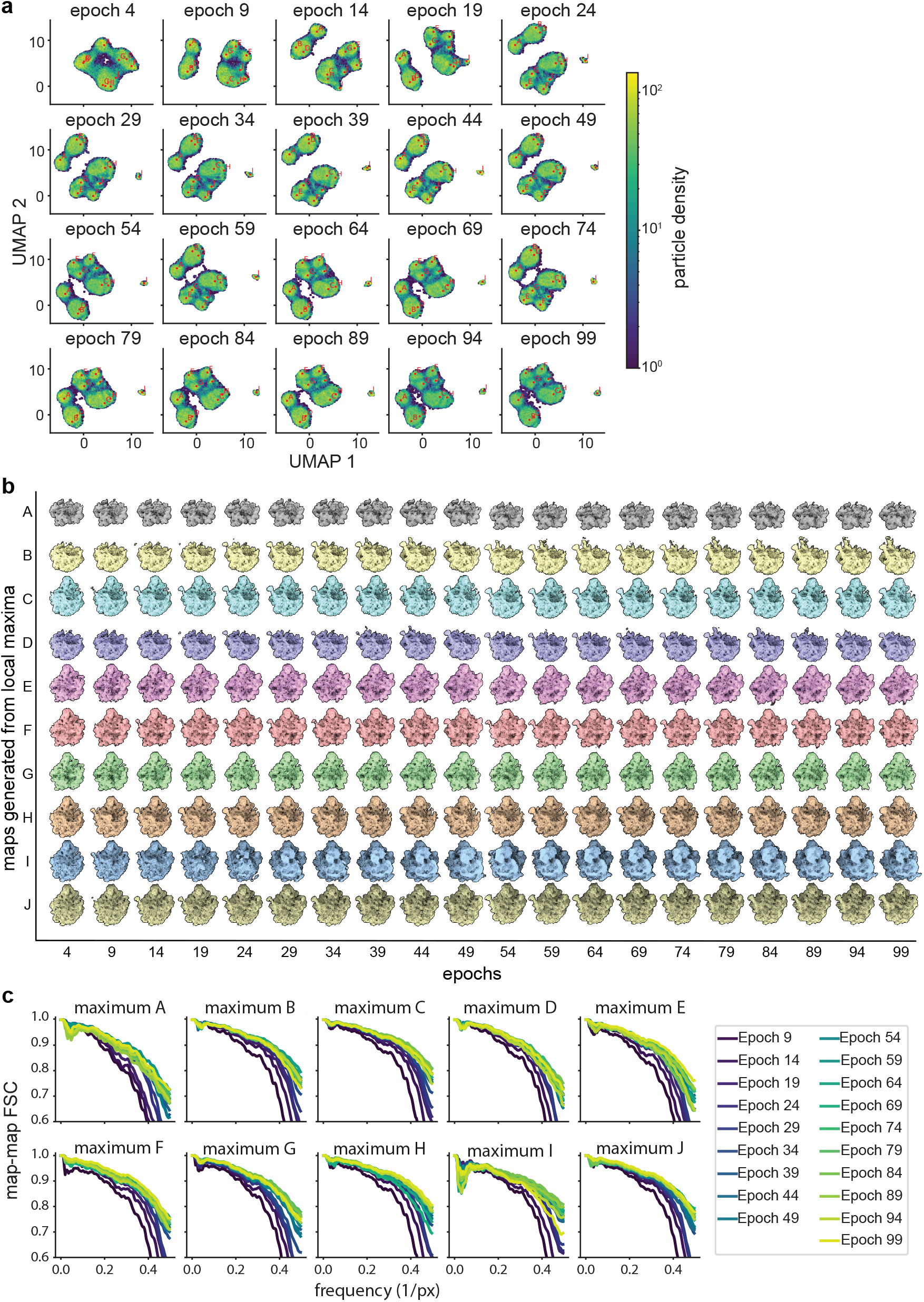
Assessing convergence of representative cryoDRGN density maps during high-resolution training. **a)** Particle sets A-J identified by the *“UMAP local maxima method”* (Glossary) mapped to prior epochs as illustrated in Extended Data Figure 2. **b)** Corresponding volumes generated from labeled positions in panel a. Note that the volumes’ gross morphology stabilizes by epoch 19-29, though maximum I stabilizes as a 70S ribosome around epoch 39. **c)** FSC plots between volumes from each local maximum offset by 5 epochs of training, as in Extended Data Figure 2. The map-to-map FSC stabilizes by epoch 39.

**Extended Data Figure 6.**
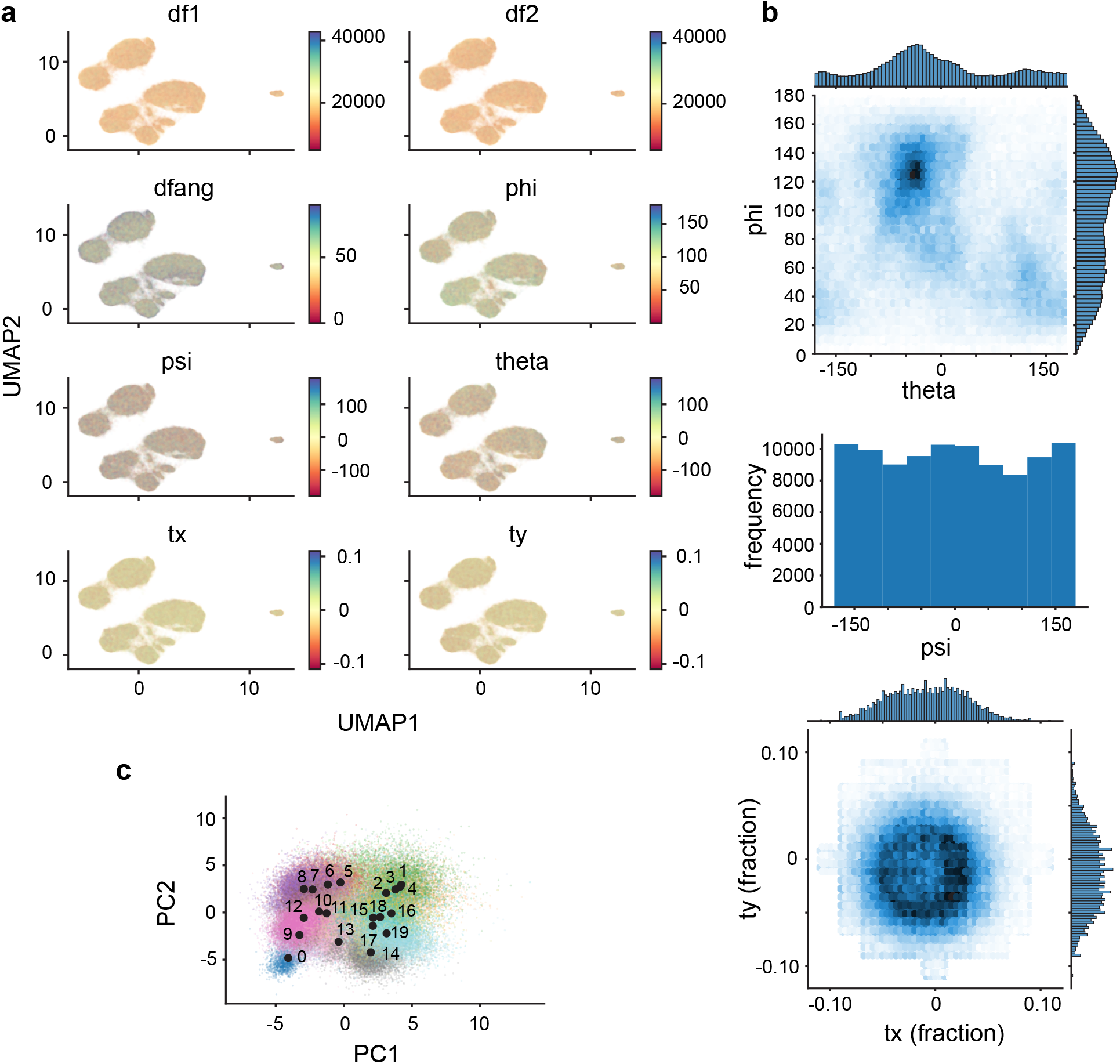
Assessing results of high-resolution training. **a)** The UMAP representation of the latent space resulting from 50 epochs of high-resolution training, colored by indicated imaging parameters. **b)** Angular and translational pose distributions. **c)** PCA of the latent space, colored by the 20 *k*-means cluster centers automatically generated by cryodrgn analyze. Numbered black dots indicate the locations in latent space of each *k*-means cluster center volume.

**Extended Data Figure 7.**
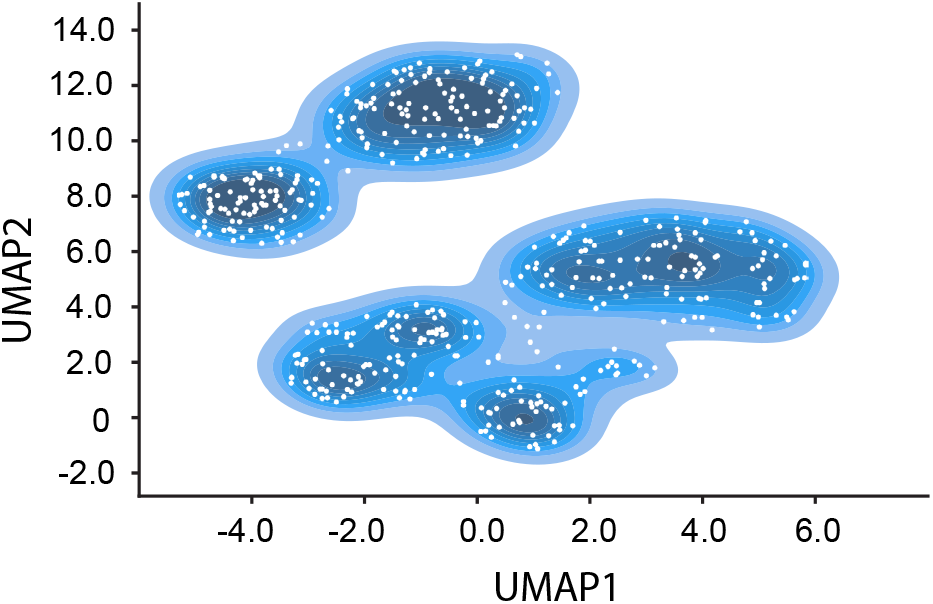
Sampled points from latent space used in subunit occupancy analysis. UMAP representation of the latent space resulting from 50 epochs of high-resolution training with contours colored with darker blues as particle density increases. Sampled points correspond to the centers of 500 *k*-means clusters and are indicated with white circles.

### Visualize data-supported structural transitions [30 minutes]

34. Return to the `cryoDRGN_viz.ipynb` analysis notebook to generate on-data centroid volumes for each class. Here, we provide the indices we define as the centroid of each of our classes in **Supplementary Table 2**. See **Supplementary Protocol 6** for detailed instructions on how to define these indices independently.
35. Having identified representative centroid indices for the varying assembly states of the 50S ribosome (**Extended Data Figure 8**), we can generate an on-data graph traversal using these points as anchors and the cryodrgn graph_traversal command. The graph traversal we highlight here showcases the B→D1→D2→D3→D4→E3→E5 assembly pathway described in Davis et al. (**Extended Data Figure 9, Supplementary Movie 3**). Note that the italicized anchor indices corresponding to the cluster centroids will vary run-to-run and should be calculated using your data as described in step 34.

~~~
cryodrgn graph_traversal 02_256_8D_1024/z.49.pkl --anchors *89122 37896 53298 81097 66910 95314 73537 51189 51011* -o 02_256_8D_1024/ analyze.49/path01.txt --out-z 02_256_8D_1024/analyze.49/z.path01.txt
cryodrgn eval_vol 02_256_8D_1024/weights.49.pkl -c 02_256_8D_1024/ config.pkl --flip --zfile 02_256_8D_1024/analyze.49/z.path01.txt -o 02_256_8D_1024/analyze.49/path01
~~~

### Validating minor states with traditional tools [1 hour]

36. While the coarse clustering described above is useful for surveying the broad landscape of structural heterogeneity within the dataset, it may obscure interesting intra-class variation. It is therefore useful to check each class individually for low-population states that differ from the rest of the class. For example, in this dataset we observe a set of volumes in class 11 with high H68 occupancy and low central protuberance occupancy. These particles correspond to the C4 class of particles identified previously by our group using cryoDRGN (Zhong et al., 2020) but which were overlooked using traditional 3D classification approaches (Davis et al., 2016).
37. Using the Jupyter notebook generated by `cryodrgn analyze`, you can extract particles corresponding to structural states of interest to conduct homogeneous refinement with tools such as cryoSPARC and RELION. Here, we select particles belonging to the C4 class *k*-means clusters, which are represented by maps 270, 283, 284, 285, and 286 in our analysis (**Extended Data Figure 10**). The map indices will vary from run to run; users should determine which particles belong to class C4 for their run by looking for maps with high H68 occupancy and low occupancy of the central protuberance block.

~~~
df = pd.read_csv (‘kmeans500_df.csv’, index_col = 0)
c4 = [270, 283, 284, 285, 286]
df_c4 = df[df[‘Kmeans500’].isin(c4)]
~~~ Having defined our selection, we can now set `ind_selected = df_c4.index` in the *“Save the index selection”* cell, and run this cell to save a .pkl file with the indices of these particles. This .pkl file can be used to filter the original .star file for import into cryoSPARC or RELION with the `cryodrgn write_star` script, as described above during particle filtering (**Figure 5D**). This concludes a preliminary cryoDRGN analysis, however users may wish to continue exploring their data using the tools we’ve described, and additional functionalities within these notebooks that are beyond the scope of this protocol. We encourage users to embrace the iterative, interactive approach to cryoDRGN analyses described herein, and hope users will find these tools valuable as they develop testable hypotheses aimed at understanding dynamic macromolecular complexes.

**Extended Data Figure 8.**
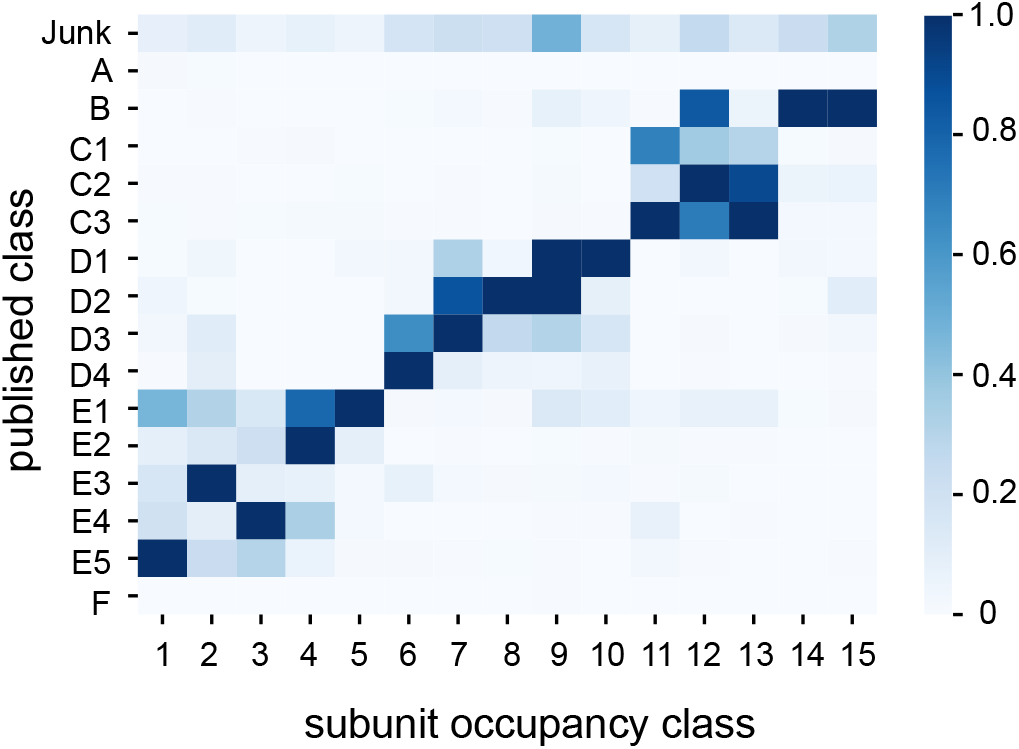
Confusion matrix of published class labels and classes assigned by subunit occupancy analysis. *K*-means 500 cluster center maps were assigned to 15 classes by subunit occupancy analysis. Particles within a given *k*-means 500 cluster are assigned to the same subunit occupancy class as the center map. Published particle labels were drawn from Davis et al. and the fractional correspondence is plotted as a heat map. Note that published classes A and F corresponded to 70S and 30S particles, respectively.

**Extended Data Figure 9.**
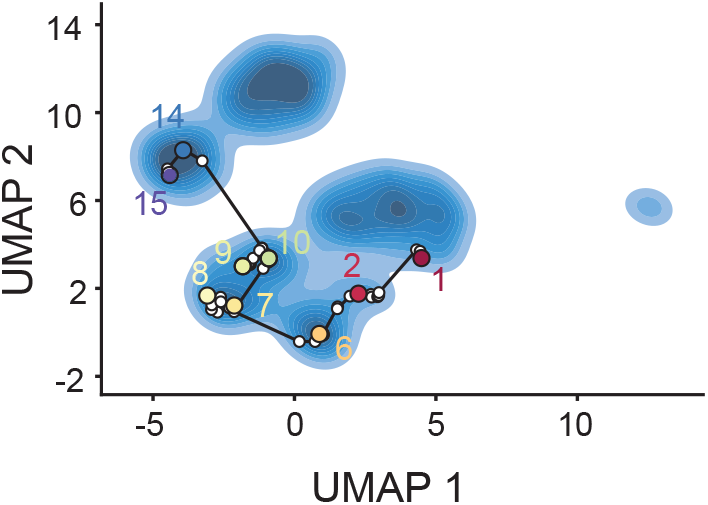
Graph traversal through latent space for the B→D1→D2→D3→D4→E3→E5 assembly pathway. Centroid volumes from the subunit occupancy classes were aligned and compared to the previously published assembly intermediate structures16 to determine approximate equivalences between published classes and subunit occupancy classes. The volumes corresponding to intermediates B, D1, D2, D3, D4, E3, and E5 were provided to `cryodrgn graph_traversal` as anchor points; the resulting path through latent space is shown. Non-anchor points are indicated with white circles, whereas anchor points and their corresponding class ID are shown with colored circles. Volumes from the complete graph traversal are shown in Supplementary Movie 3.

**Extended Data Figure 10.**
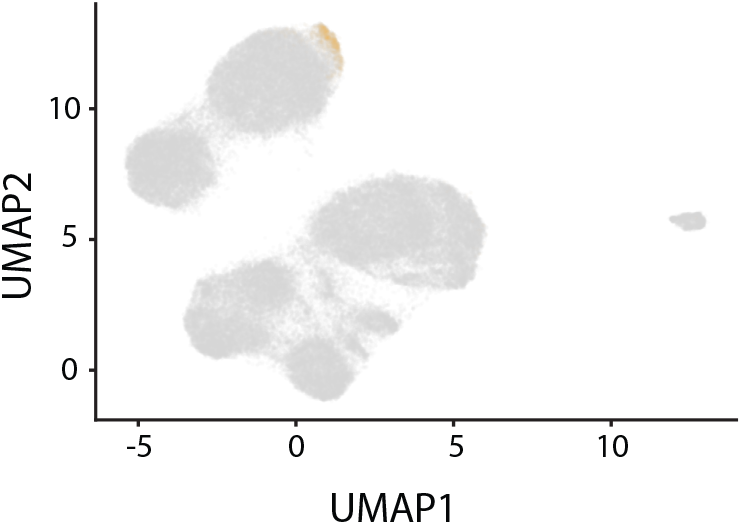
Selection of particles corresponding to the C4 minor class. Particles (1,149) in the C4 class were identified by subunit occupancy analysis and are highlighted in orange.

## TROUBLESHOOTING

| Step | Problem | Possible reason | Possible solution |
| --- | --- | --- | --- |
| 4 | Out of memory error when running `cryodrgn downsample` | The particle stack is too large to fit into memory | Add the `--chunk` flag to the `cryodrgn downsample` command |
| 5 | Back projection is noisy, discontinuous, or does not resemble consensus refinement | Incorrect pose and CTF metadata were supplied, or pose and CTF parameters were incorrectly mapped to particles. Noisy maps may also result from using a small number of particles used in the back projection (default: 10,000). Users may also have not applied the correct `--uninvert-data` convention, which determines whether the data is light-on-dark or dark-on-light | Verify that the correct pose and CTF parameters were supplied during parsing and that the particle stack originated from, and contains the same particle index/order as the pose and CTF parameter metadata. If the volume is very noisy, re-run `cryodrgn backproject_voxel` with a larger number of particles using `--first` flag. Check whether the correct `--uninvert-data` convention for the dataset is followed by running `cryodrgn backproject_voxel` with and without `--uninvert-data` |
| 6, 18 | Out of memory error shortly after starting `cryodrgn train_vae` | The particle stack is too large to preload into memory | Append `--lazy` to the `cryodrgn train_vae` command to allow on-the-fly image loading, further downsample particles, or train on a subset of the particle stack |
| 6, 18 | CUDA out of memory error during `cryodrgn train_vae` | Batch size may be set too large for your GPU's memory capacity | Manually decrease batchsize with the `--batchsize` flag in the `cryodrgn train_vae` command or further downsample particle images |
| 6, 18 | Assertion error during `cryodrgn train_vae` similar to `assert (coords[... , 0:3].abs() - 0.5 < 1e-4).all()` | Infrequent issues with numerical instability using `--amp` may cause this assertion to fail | Restart `cryodrgn train_vae` without `--amp` |
| 7, 9, 19, 22-23 | Volumes generated after training appear non-continuous or hollow in the center of the box | Users failed to apply correct `--uninvert-data` flag | Run `cryodrgn backproject_voxel` with and without the `--uninvert-data` flag and determine which convention is applicable, then re-run `cryodrgn train_vae` as necessary |
| 7, 9, 19, 22-23 | Volumes generated after training all resemble "junk" | Volumes may be displayed at too permissive an isosurface. Alternatively, data may have been parsed incorrectly in the preprocessing steps | Increase the isosurface threshold for display. Run `cryodrgn backproject_voxel` to determine whether poses and CTF parameters were correctly parsed. |
| 7, 9, 19, 22-23 | Volumes generated after training all appear homogeneous | For datasets other than EMPIAR-10076, this may be caused by too much upstream filtering prior to cryoDRGN training | Restart cryoDRGN training with an unfiltered dataset. |
| All | Jupyter notebooks aren't behaving as described in the protocol | Cells may have been run out of order or may reference outdated variables | Restart the kernel and run the notebook again in order from top to bottom |

## TIMING

The required time to run this protocol is dependent on the hardware users have available. We provide approximate timings in each section of the protocol based on our hardware described in **Materials**. For users who seek to employ this protocol on their own datasets, the primary determinant of the timing will be how long the cryoDRGN model training steps require, as these steps are the most expensive in terms of both time and computational resources. We generally recommend training *“high-resolution”* cryoDRGN models at a boxsize of 256 pixels, as computational time can become prohibitive with significantly larger boxes. For very large datasets or datasets with large boxsizes, users may find it useful to employ the `cryodrgn preprocess` command instead of `cryodrgn downsample`, as this command changes some of the preprocessing steps to minimize downstream memory usage and obviates the need for using on-the-fly image loading via the `--lazy` flag, which significantly increases training times. Instructions for how to use `cryodrgn preprocess` are available at https://github.com/zhonge/cryodrgn.

## ANTICIPATED RESULTS

This protocol describes the training of a cryoDRGN model on a highly heterogeneous exemplar dataset (EMPIAR-10076), as well as the systematic characterization of the resulting structural ensemble. Following the protocol, users produce the following principal outputs:

1. A latent embedding for each particle in the input stack.
2. A decoder network able to generate an arbitrary number of volumes from embeddings across latent space. This decoder network can then be used, as shown in this protocol, to explore the structural landscape of the dataset by sampling the 3D volumes found in different positions of latent space.
3. A representative ensemble of volumes sampled from across latent space using the decoder network, which can be directly visualized and used for downstream landscape analysis.
4. A matrix of occupancy values for each structural element in each sampled volume, which can be clustered and represented as a heatmap, and which can be used for quantitative analysis of the sample’s structural heterogeneity.

Though the precise nature of the heterogeneity uncovered is dataset-dependent, and aspects of the analysis – notably how clustered or featured the distribution of latent embeddings is – may differ from the analysis of this example dataset, users should be able to follow this protocol on their own datasets to produce a similar set of outputs.

## DATA AVAILABILITY

All final and intermediate results presented in this protocol are available at https://doi.org/10.5281/zenodo.5164127

## CODE AVAILABILITY

The software and scripts used in these analyses are available at https://github.com/zhonge/cryodrgn (version 0.3.5) and https://github.com/lkinman/occupancy-analysis (version 0.1.2), as described in **Materials**. All code is available through the open source GPL-3.0 License.

## AUTHOR CONTRIBUTIONS

Conceptualization - All; Funding acquisition - BB, JHD; Investigation - LK, BMP, EDZ, JHD; Software - LK, BMP, EDZ, JHD; Supervision - BB, JHD; Visualization - LK, BMP, JHD; Writing [original draft] - LK, BMP, EDZ; Writing [editing] – LK, BMP, EDZ, JHD.

## ACKNOWLEDGMENTS

We thank the MIT-IBM Satori team for GPU computing resources and support. This work was funded by the NSF GRFP Fellowship to E.D. Z., NIH grant R01-GM081871 to B.B., NSFCAREER-2046778 and NIH grant R01-GM144542 to J.H.D., and a grant from the MIT J-Clinic for Machine Learning and Health to J.H.D. and B.B. Research in the Davis lab is supported by the Alfred P. Sloan Foundation, the James H. Ferry Fund, and the Whitehead Family.

## SUPPLEMENTAL PROTOCOLS

### Supplementary protocol 1. Installing cryoDRGN version 0.3.5

1. Instructions for installing the latest version of cryoDRGN are available at https://github.com/zhonge/cryodrgn. For consistency with our results, we recommend using version 0.3.5 that we employed in this protocol. It can be installed using github as described below. To set up the conda environment, run the following commands:

~~~
conda create --name cryodrgn python=3.7
conda activate cryodrgn
conda install pytorch cudatoolkit=10.2 -c pytorch
conda install pandas seaborn scikit-learn
conda install umap-learn jupyterlab ipywidgets cufflinks-py “nodejs>=15.12.0” -c conda-forge
conda update typing_extensions -c conda-forge
jupyter labextension install @jupyter-widgets/jupyterlab-manager --no-build
jupyter labextension install jupyterlab-plotly --no-build jupyter labextension install plotlywidget --no-build jupyter lab build
~~~ **Critical:** Ensure that you install cudatoolkit and pytorch versions compatible with your graphics card and drivers. For example, your CUDA version is returned by the command nvidia-smi, and generally the latest pytorch version (built for your CUDA version and python 3.7) will be appropriate. See pytorch.org for more details on how to install pytorch.
2. Optionally install NVIDIA’s Apex library to enable `--amp` acceleration via the following commands:

~~~
git clone https://github.com/NVIDIA/apex
cd apex
pip install -v --disable-pip-version-check --no-cache-dir./
~~~
3. Optionally install the CUDA machine learning library for faster UMAP embeddings in `analyze_ convergence.py`.

~~~
conda install cuml -c rapidsai-nightly -c rapidsai -c nvidia -c conda-forge
~~~
4. Clone version 0.3.5 from GitHub:

~~~
git clone https://github.com/zhonge/cryodrgn.git
cd cryodrgn
git checkout tags/0.3.5
python setup.py -q install
~~~

### Supplementary protocol 2. Creating a consensus refinement in cryoSPARC

1. Run an import particle stack job by specifying `L17Combine_weight_local.mrcs` as the particle data path and `Parameters.star` as the particle meta path. Note that the data sign needs to be flipped to dark-on-light.
2. Run an *ab initio* reconstruction job with default parameters.
3. Run a homogeneous refinement job with default parameters. Note that we generally suggest performing reconstructions without imposed symmetry (*i.e.* C1) as it preserves potentially interesting heterogeneity.
4. Copy the refined particles.cs file, whose name should resemble `cryosparc_P4_J33_004_ particles.cs`, to the working cryoDRGN directory where the full dataset is stored.

### Supplementary protocol 3. Setting up port forwarding via SSH

1. SSH port forwarding can be set up at the time of login using the following command and replacing `remote_username` and `remote_host_name` with the appropriate values:

~~~
ssh -N -f -L localhost:8888:localhost:8888 remote_username@remote_host_name
~~~ If you are running your jupyter notebook on a worker node in a compute cluster, as opposed to a local workstation, we suggest the following alternative port forwarding command:

~~~
ssh -t -t username@cluster-head-node -L 8888:localhost:8888 ssh active-worker-node -L 8888:localhost:8888
~~~
2. To open a Jupyter notebook, enter the command `jupyter lab --no-browser --port 8888` into the terminal, and navigate to localhost:8888 in a web browser on your local computer.

### Supplementary protocol 4. Generating segmented PDB chains for subunit occupancy analysis

1. Open PyMOL and use the command-line interface to retrieve an atomic model of the 70S ribosome from the PDB: `fetch 4ybb`
2. Delete atoms outside the region of interest. For example, to generate the segmented .pdb of the 5S rRNA, we use:

~~~
sele not_5s, not chain CB
~~~ Then select “Remove atoms’ from the drop-down “Action” menu in the not_5s selection. This will delete all non-5S atoms.
3. Segment the map into chains if necessary. If you want to do occupancy analysis on whole protein subunits, this is likely unnecessary, as the chains are likely already defined in the atomic model. If you want to define your own subunits for occupancy analysis as we do here, you can do so using the alter command as shown below, again for the examples of the 5S rRNA:

~~~
alter (resi 1-14,108-120), chain=‘A’
alter (resi 15-27,60-68), chain=‘B’
alter (resi 28-59), chain=‘C’
alter (resi 78-99), chain=‘D’
alter (resi 69-77,100-107), chain=‘E’
~~~
4. After you have made all the chain alterations, save the .pdb file with a new name, e.g. `RNA_5S.pdb` using the “Export Molecule” command. Note that to create more than 26 chains, you will need to use multiple .pdb files, each containing at most 26 chain IDs.

### Supplementary protocol 5. Aligning segmented PDB models for subunit occupancy analysis

1. The.pdb files must now be aligned to your cryoDRGN sampled maps. Open one of the generated 500 maps (*e.g.* vol_000.mrc) in ChimeraX. Aim to select a map that has high occupancy of most elements of your structure to ensure a good alignment. Because maps with adjacent indices (*e.g.*vol_000 and vol_001) are often structurally similar as they are sampled from proximal locations in latent space, users are advised to find a mature map by downloading 20 random volumes from the set of 500.
2. Open all the .pdb files (`prots1.pdb, prots2.pdb, RNA_5S.pdb, RNA1.pdb, RNA2. pdb, RNA3.pdb, RNA4.pdb`). These should now be models #2-8 in your ChimeraX session.
3. Select models #2-8 with the command select #2-8. Provide a rough manual alignment between the selected atoms and the example map, using the “Rotate model” and “Move model” right mouse modes.
4. Having provided a rough manual alignment, use the Tools > Volume Data > Fit in Map option to fit your .pdb files in the map. Choose to fit “selected atoms” in your example map, making sure that all the .pdb model files are still selected.
5. Save each of the .pdb files individually using File > Save, and selecting .pdb file type. Be sure you have the correct .pdb model selected in the Models selection box, and that you select the option to “Save relative to model:”, with the example map selected as the model.

### Supplementary protocol 6. Identifying centroid volumes for subunit occupancy volume classes

1. Use pandas to load dataframe you saved with information about which *k*-means 500 class each particle corresponds to.

~~~
df = pd.read_csv(‘kmeans500_df.csv’, index_col = 0)
~~~
2. To save the volume classes defined by clustering in the `occupancy_analysis.ipynb` Jupyter notebook, run the cells in the *“Extract classes from clustering”* section. This will save the class assignments as a .pkl file that you can load into the `cryoDRGN_viz.ipynb` notebook.
3. Open the volume class assignments .pkl file in the `cryoDRGN_viz.ipynb` notebook, changing the name or relative path of the .pkl file name as necessary in the code below.

~~~
classes = utils.load_pkl(‘../../vol class.pkl’)
~~~
4. Identify the nearest on-data point to the median z-coordinates of each class. The resulting variable `nearest_inds` contains the indices in your dataframe of the centroid particles. You can then generate volumes at these indices as before using the volume generation cells of the Jupyter notebook.

~~~
median_coords = np.empty([len(classes.keys()), z.shape[1]], dtype =‘float64’)
z_list = df.columns[df.columns.str.contains(‘z’)]
for i in classes.keys():
   df.loc[df[df[‘Kmeans500’].isin(classes[i])].index, ‘volume_class’] = i
   sub = df[df[‘volume_class’] == i][z_list]
   median coords[i, :] = np.array(sub.median(axis = 0))
df z = df[z_list]
neighbor_dists = pd.DataFrame(distance.cdist(median_coords, df_z, ‘euclidean’))
nearest_inds = neighbor_dists.idxmin(axis = 1)
~~~

## SUPPLEMENTARY TABLES

**Supplementary Table 1:** Residue and chain assignment for subunit occupancy analysis. Ribosomal RNA helices and ribosomal proteins are numbered as in Davis et al.

| Subunit | PDB ID | PDB Chain | Residues | Segmented file name | Segmented file chain |
| --- | --- | --- | --- | --- | --- |
| H1 | 4YBB | CA | 1-12,2895-2904 | RNA1.pdb | A |
| H2 | 4YBB | CA | 13-30,510-531 | RNA1.pdb | B |
| H3 | 4YBB | CA | 31-32,473-474 | RNA1.pdb | C |
| H4 | 4YBB | CA | 33-47,431-451 | RNA1.pdb | D |
| H5 | 4YBB | CA | 48-56,114-120 | RNA1.pdb | E |
| H6 | 4YBB | CA | 57-74 | RNA1.pdb | F |
| H7 | 4YBB | CA | 75-113 | RNA1.pdb | G |
| H8 | 4YBB | CA | 121-130 | RNA1.pdb | H |
| H9 | 4YBB | CA | 131-148 | RNA1.pdb | I |
| H10 | 4YBB | CA | 147-177 | RNA1.pdb | J |
| H11 | 4YBB | CA | 178-218,319-323 | RNA1.pdb | K |
| H12 | 4YBB | CA | 219-232 | RNA1.pdb | L |
| H13 | 4YBB | CA | 233-262 | RNA1.pdb | M |
| H14 | 4YBB | CA | 263-269,424-430 | RNA1.pdb | N |
| H16 | 4YBB | CA | 269-280,360-370 | RNA1.pdb | O |
| H18 | 4YBB | CA | 281-298,340-359 | RNA1.pdb | P |
| H19 | 4YBB | CA | 299-318 | RNA1.pdb | Q |
| H20 | 4YBB | CA | 324-339 | RNA1.pdb | R |
| H21 | 4YBB | CA | 371-404 | RNA1.pdb | S |
| H22 | 4YBB | CA | 405-423 | RNA1.pdb | T |
| H23 | 4YBB | CA | 452-472 | RNA1.pdb | U |
| H24 | 4YBB | CA | 475-509 | RNA1.pdb | V |
| H25 | 4YBB | CA | 532-561 | RNA1.pdb | W |
| H25a | 4YBB | CA | 562-578 | RNA1.pdb | X |
| H26 | 4YBB | CA | 579-586,1251-1261 | RNA1.pdb | Y |
| H27 | 4YBB | CA | 587-602,655-670 | RNA1.pdb | Z |
| H28 | 4YBB | CA | 603-625 | RNA2.pdb | A |
| H29 | 4YBB | CA | 626-636 | RNA2.pdb | B |
| H31 | 4YBB | CA | 637-654 | RNA2.pdb | C |
| H32 | 4YBB | CA | 671-683,790-809 | RNA2.pdb | D |
| H33 | 4YBB | CA | 684-698,763-775 | RNA2.pdb | E |
| H34 | 4YBB | CA | 699-733 | RNA2.pdb | F |
| H35 | 4YBB | CA | 734-762 | RNA2.pdb | G |
| H35a | 4YBB | CA | 776-789 | RNA2.pdb | H |
| H36 | 4YBB | CA | 810-821,1186-1195 | RNA2.pdb | I |
| H37 | 4YBB | CA | 822-835 | RNA2.pdb | J |
| H38 | 4YBB | CA | 836-942 | RNA2.pdb | K |
| H39 | 4YBB | CA | 943-973 | RNA2.pdb | L |
| H40 | 4YBB | CA | 974-990 | RNA2.pdb | M |
| H41 | 4YBB | CA | 991-1025,1133-1163 | RNA2.pdb | N |
| H42 | 4YBB | CA | 1026-1056,1103-1132 | RNA2.pdb | O |
| H43 | 4YBB | CA | 1057-1081 | RNA2.pdb | P |
| H44 | 4YBB | CA | 1087-1102 | RNA2.pdb | Q |
| H45 | 4YBB | CA | 1164-1185 | RNA2.pdb | R |
| H46 | 4YBB | CA | 1196-1250 | RNA2.pdb | S |
| H26a | 4YBB | CA | 1262-1270,2010-2017 | RNA2.pdb | T |
| H47 | 4YBB | CA | 1271-1294 | RNA2.pdb | U |
| H48 | 4YBB | CA | 1295-1302,1640-1647 | RNA2.pdb | V |
| H49 | 4YBB | CA | 1303-1306,1622-1625 | RNA2.pdb | W |
| H49b | 4YBB | CA | 1307-1313,1603-1608 | RNA2.pdb | X |
| H50 | 4YBB | CA | 1314-1338 | RNA2.pdb | Y |
| H51 | 4YBB | CA | 1339-1347,1599-1602 | RNA2.pdb | Z |
| H52 | 4YBB | CA | 1348-1382 | RNA3.pdb | A |
| H53 | 4YBB | CA | 1383-1404 | RNA3.pdb | B |
| H54 | 4YBB | CA | 1405-1417,1581-1598 | RNA3.pdb | C |
| H55 | 4YBB | CA | 1418-1428,1569-1580 | RNA3.pdb | D |
| H49a | 4YBB | CA | 1609-1621 | RNA3.pdb | E |
| H56 | 4YBB | CA | 1429-1444,1547-1564 | RNA3.pdb | F |
| H57 | 4YBB | CA | 1445-1466 | RNA3.pdb | G |
| H58 | 4YBB | CA | 1467-1525 | RNA3.pdb | H |
| H59 | 4YBB | CA | 1526-1546 | RNA3.pdb | I |
| H60 | 4YBB | CA | 1626-1639 | RNA3.pdb | J |
| H61 | 4YBB | CA | 1648-1678,1990-2009 | RNA3.pdb | K |
| H62 | 4YBB | CA | 1679-1706 | RNA3.pdb | L |
| H63 | 4YBB | CA | 1707-1751 | RNA3.pdb | M |
| H64 | 4YBB | CA | 1758-1773,1977-1989 | RNA3.pdb | N |
| H65 | 4YBB | CA | 1774-1790 | RNA3.pdb | O |
| H66 | 4YBB | CA | 1791-1828 | RNA3.pdb | P |
| H67 | 4YBB | CA | 1829-1834,1970-1976 | RNA3.pdb | Q |
| H68 | 4YBB | CA | 1835-1905 | RNA3.pdb | R |
| H69 | 4YBB | CA | 1906-1924 | RNA3.pdb | S |
| H71 | 4YBB | CA | 1932-1969 | RNA3.pdb | T |
| H72 | 4YBB | CA | 2018-2042 | RNA3.pdb | U |
| H73 | 4YBB | CA | 2043-2057,2611-2625 | RNA3.pdb | V |
| H74 | 4YBB | CA | 2058-2074,2430-2451 | RNA3.pdb | W |
| H75 | 4YBB | CA | 2075-2092,2226-2245 | RNA3.pdb | X |
| H76 | 4YBB | CA | 2093-2114,2179-2196 | RNA3.pdb | Y |
| H77 | 4YBB | CA | 2115-2126,2169-2178 | RNA3.pdb | Z |
| H78 | 4YBB | CA | 2127-2168 | RNA4.pdb | A |
| H79 | 4YBB | CA | 2197-2225 | RNA4.pdb | B |
| H80 | 4YBB | CA | 2246-2258 | RNA4.pdb | C |
| H81 | 4YBB | CA | 2259-2281 | RNA4.pdb | D |
| H82 | 4YBB | CA | 2282-2286,2382-2390 | RNA4.pdb | E |
| H83 | 4YBB | CA | 2287-2296,2335-2344 | RNA4.pdb | F |
| H84 | 4YBB | CA | 2297-2321 | RNA4.pdb | G |
| H85 | 4YBB | CA | 2322-2334 | RNA4.pdb | H |
| H86 | 4YBB | CA | 2345-2371 | RNA4.pdb | I |
| H87 | 4YBB | CA | 2372-2381 | RNA4.pdb | J |
| H88 | 4YBB | CA | 2391-2429 | RNA4.pdb | K |
| H89 | 4YBB | CA | 2452-2504 | RNA4.pdb | L |
| H90 | 4YBB | CA | 2505-2517,2567-2586 | RNA4.pdb | M |
| H91 | 4YBB | CA | 2518-2546 | RNA4.pdb | N |
| H92 | 4YBB | CA | 2547-2561 | RNA4.pdb | O |
| H93 | 4YBB | CA | 2587-2610 | RNA4.pdb | P |
| H94 | 4YBB | CA | 2626-2643,2771-2788 | RNA4.pdb | Q |
| H95 | 4YBB | CA | 2644-2675 | RNA4.pdb | R |
| H96 | 4YBB | CA | 2676-2731 | RNA4.pdb | S |
| H97 | 4YBB | CA | 2732-2770 | RNA4.pdb | T |
| H98 | 4YBB | CA | 2789-2805 | RNA4.pdb | U |
| H99 | 4YBB | CA | 2806-2814,2886-2894 | RNA4.pdb | V |
| H100 | 4YBB | CA | 2815-2831 | RNA4.pdb | W |
| H101 | 4YBB | CA | 2832-2885 | RNA4.pdb | X |
| H1_5S | 4YBB | CB | 1-14,108-120 | RNA_5S.pdb | A |
| H2_5S | 4YBB | CB | 15-27,60-68 | RNA_5S.pdb | B |
| H3_5S | 4YBB | CB | 28-59 | RNA_5S.pdb | C |
| H4_5S | 4YBB | CB | 78-99 | RNA_5S.pdb | D |
| H5_5S | 4YBB | CB | 69-77,100-107 | RNA_5S.pdb | E |
| uL2 | 4YBB | CC | all | prots1.pdb | A |
| uL3 | 4YBB | CD | all | prots1.pdb | B |
| uL4 | 4YBB | CE | all | prots1.pdb | C |
| uL5 | 4YBB | CF | all | prots1.pdb | D |
| uL6 | 4YBB | CG | all | prots1.pdb | E |
| bL9 | 4YBB | CH | all | prots1.pdb | F |
| uL11 | 4YBB | CJ | all | prots1.pdb | G |
| uL13 | 4YBB | CK | all | prots1.pdb | H |
| uL14 | 4YBB | CL | all | prots1.pdb | I |
| uL15 | 4YBB | CM | all | prots1.pdb | J |
| uL16 | 4YBB | CN | all | prots1.pdb | K |
| bL17 | 4YBB | CO | all | prots1.pdb | L |
| uL18 | 4YBB | CP | all | prots1.pdb | M |
| bL19 | 4YBB | CQ | all | prots1.pdb | N |
| bL20 | 4YBB | CR | all | prots1.pdb | O |
| bL21 | 4YBB | CS | all | prots1.pdb | P |
| uL22 | 4YBB | CT | all | prots1.pdb | Q |
| uL23 | 4YBB | CU | all | prots1.pdb | R |
| uL24 | 4YBB | CV | all | prots1.pdb | S |
| bL25 | 4YBB | CW | all | prots1.pdb | T |
| bL27 | 4YBB | CX | all | prots1.pdb | U |
| bL28 | 4YBB | CY | all | prots1.pdb | V |
| uL29 | 4YBB | CZ | all | prots1.pdb | W |
| uL30 | 4YBB | C0 | all | prots1.pdb | X |
| bL32 | 4YBB | C1 | all | prots1.pdb | Y |
| bL33 | 4YBB | C2 | all | prots1.pdb | Z |
| bL34 | 4YBB | C3 | all | prots2.pdb | A |
| bL35 | 4YBB | C4 | all | prots2.pdb | B |
| bL36 | 4YBB | C5 | all | prots2.pdb | C |

**Supplementary Table 2:** Particle stack indices for centroid volumes of subunit occupancy classes. Note that these indices are only relevant for the provided pre-computed results and users should select alternative indices when training new cryoDRGN models.

| Class | Centroid index |
| --- | --- |
| 1 | 51011 |
| 2 | 51189 |
| 3 | 80371 |
| 4 | 9177 |
| 5 | 74182 |
| 6 | 73537 |
| 7 | 95314 |
| 8 | 66910 |
| 9 | 53298 |
| 10 | 81097 |
| 11 | 11144 |
| 12 | 71755 |
| 13 | 46961 |
| 14 | 37896 |
| 15 | 89122 |

## SUPPLEMENTARY MOVIES

**Supplementary Movie 1. PC1 trajectory from high resolution training.**
Density maps sampled along PC1 were automatically generated by the `cryodrgn analyze` command. Volumes are displayed at the same isosurface level, and the movie progresses from low to high PC1 value strictly along the PC1 axis.

**Supplementary Movie 2. PC2 trajectory from high resolution training.**
Density maps sampled along PC2 were automatically generated by the `cryodrgn analyze` command. Volumes are displayed at the same isosurface level, and the movie progresses from low to high PC2 value strictly along the PC2 axis.

**Supplementary Movie 3. Graph traversal of the B**→**D1**→**D2**→**D3**→**D4**→**E3**→**E5 assembly pathway.**
Graph traversal pathway was generated using the `cryodrgn graph_traversal` command as described in the protocol. The path taken by the traversal through latent space is shown in **Extended Data Figure 9**. All volumes are displayed at the same isosurface level.

## Footnotes

1 Equivalent commands exist for preprocessing poses and CTF from RELION outputs as `parse_pose_star` and `parse_ctf_star`. If using a star file from RELION 3.1 or later, include the `--relion31` flag in this command.

## REFERENCES

Cheng, Y., Grigorieff, N., Penczek, P. A. & Walz, T. A primer to single-particle cryo-electron microscopy. Cell 161, 438–449, doi:10.1016/j.cell.2015.03.050 (2015).

Dashti, A. et al. Trajectories of the ribosome as a Brownian nanomachine. Proc Natl Acad Sci U S A 111, 17492–17497, doi:10.1073/pnas.1419276111 (2014).

Dashti, A. et al. Retrieving functional pathways of biomolecules from single-particle snapshots. Nat Commun 11, 4734, doi:10.1038/s41467-020-18403-x (2020).

Davis, J. H. et al. Modular Assembly of the Bacterial Large Ribosomal Subunit. Cell 167, 1610–1622 e1615, doi:10.1016/j.cell.2016.11.020 (2016).

Davis, J. H. & Williamson, J. R. Structure and dynamics of bacterial ribosome biogenesis. Philos Trans Soc B 372, doi:10.1098/rstb.2016.0181 (2017).

Goddard, T. D. et al. UCSF ChimeraX: Meeting modern challenges in visualization and analysis. Protein Sci 27, 14–25, doi:10.1002/pro.3235 (2018).

Gui, M. et al. Structures of radial spokes and associated complexes important for ciliary motility. Nat Struct Mol Biol 28, 29–37, doi:10.1038/s41594-020-00530-0 (2021).

Grant, T., Rohou, A. & Grigorieff, N. cisTEM, user-friendly software for single-particle image processing. Elife 7, doi:10.7554/eLife.35383 (2018).

Haselbach, D. et al. Long-range allosteric regulation of the human 26S proteasome by 20S proteasome-target-ing cancer drugs. Nat Commun 8, 15578, doi:10.1038/ncomms15578 (2017).

Kingma, D. & Welling, M. Auto-encoding variational bayes. 2nd International Conference on Learning Representations (2013).

Ludtke, S. & Chen, M. Deep learning based mixed-dimensional GMM for characterizing variability in CryoEM. arXiv, doi:arXiv:2101.10356 (2021).

Lyumkis, D. Challenges and opportunities in cryo-EM single-particle analysis. J Biol Chem 294, 5181–5197, doi:10.1074/jbc.REV118.005602 (2019).

Nakane, T., Kimanius, D., Lindahl, E. & Scheres, S. H. Characterisation of molecular motions in cryo-EM sin-gle-particle data by multi-body refinement in RELION. eLife 7, doi:10.7554/eLife.36861 (2018).

McInnes, L., Healy, J. & Melville, J. UMAP: Uniform Manifold Approximation and Projection for dimension reduction. doi:https://arxiv.org/abs/1802.03426 (2018).

Punjani, A. & Fleet, D. J. 3D variability analysis: Resolving continuous flexibility and discrete heterogeneity from single particle cryo-EM. J Struct Biol 213, 107702, doi:10.1016/j.jsb.2021.107702 (2021).

Punjani, A. & Fleet, D. J. 3D Flexible Refinement: Structure and Motion of Flexible Proteins from Cryo-EM. bioRx-iv, 2021.2004.2022.440893, doi:10.1101/2021.04.22.440893 (2021).

Punjani, A., Rubinstein, J. L., Fleet, D. J. & Brubaker, M. A. cryoSPARC: algorithms for rapid unsupervised cryo-EM structure determination. Nat Methods 14, 290–296, doi:10.1038/nmeth.4169 (2017).

Punjani, A., Zhang, H. & Fleet, D. J. Non-uniform refinement: adaptive regularization improves single-particle cryo-EM reconstruction. Nat Methods 17, 1214–1221, doi:10.1038/s41592-020-00990-8 (2020).

Rabuck-Gibbons, J. N., Lyumkis, D. & Williamson, J. R. Quantitative Mining of Compositional Hetero-geneity in Cryo-EM Datasets of Ribosome Assembly Intermediates. bioRxiv, 2021.2006.2023.449614, doi:10.1101/2021.06.23.449614 (2021).

Rosenbaum, D. et al. Inferring a Continuous Distribution of Atom Coordinates from Cryo-EM Images using VAEs. arXiv, doi:https://arxiv.org/abs/2106.14108v1 (2021).

Scheres, S. H. RELION: implementation of a Bayesian approach to cryo-EM structure determination. J Struct Biol 180, 519–530, doi:10.1016/j.jsb.2012.09.006 (2012).

Serna, M. Hands on Methods for High Resolution Cryo-Electron Microscopy Structures of Heterogeneous Mac-romolecular Complexes. Front Mol Biosci 6, 33, doi:10.3389/fmolb.2019.00033 (2019).

Sun, J.Y., Kinman, L.K., Ortega, J. & Davis, J.H. KsgA facilitates ribosomal small subunit maturation by proofreading a key structural lesion. bioRxiv, 2022.07.13.499473, doi:https://doi.org/10.1101/2022.07.13.499473 (2022).

Trabuco, L. G., Villa, E., Schreiner, E., Harrison, C. B. & Schulten, K. Molecular dynamics flexible fitting: a practical guide to combine cryo-electron microscopy and X-ray crystallography. Methods 49, 174–180, doi:10.1016/j.ymeth.2009.04.005 (2009).

von Loeffelholz, O. et al. Focused classification and refinement in high-resolution cryo-EM structural analysis of ribosome complexes. Curr Opin Struct Biol 46, 140–148, doi:10.1016/j.sbi.2017.07.007 (2017).

Wu, M. & Lander, G. C. Present and Emerging Methodologies in Cryo-EM Single-Particle Analysis. Biophys J 119, 1281–1289, doi:10.1016/j.bpj.2020.08.027 (2020).

Zivanov, J. et al. New tools for automated high-resolution cryo-EM structure determination in RELION-3. eLife 7, doi:10.7554/eLife.42166 (2018).

Zivanov, J., Nakane, T. & Scheres, S. H. W. A Bayesian approach to beam-induced motion correction in cryo-EM single-particle analysis. IUCrJ 6, 5–17, doi:10.1107/S205225251801463X (2019).

Zhong, E., Bepler, T., Berger, B. & Davis, J. CryoDRGN: Reconstruction of Heterogeneous cryo-EM Structures Using Neural Networks. Nature Methods, doi:10.1038/s41592-020-01049-4 (2020).

Zhong, E., Bepler, T., Davis, J. & Berger, B. Reconstructing continuously heterogeneous structures from single particle cryo-EM with deep generative models. arXiv, doi:arXiv:1909.05215 (2019).

Zhong, E. D., Lerer, A., Davis, J. H. & Berger, B. Exploring generative atomic models in cryo-EM reconstruction. arXiv, doi:https://arxiv.org/abs/2107.01331v1 (2021).

